# Multi-ion permeation and dynamic conductance modulation in connexin gap junction channels

**DOI:** 10.64898/2026.08.12.744469

**Authors:** Bassam G. Haddad, Daniel M. Zuckerman, Steve L. Reichow

**Author notes:** **Correspondence:** Steve L. Reichow, Bassam G. Haddad.

## Abstract

Gap junction channels formed by connexins mediate direct intercellular communication and are essential for electrical signaling and tissue homeostasis. Despite their large, solvent-accessible pores, connexin channels exhibit distinct conductance, selectivity, and rectification properties, but the molecular mechanisms underlying these behaviors remain incompletely understood. Here, we performed ∼67 μs of all-atom computational electrophysiology simulations of connexin-46 (Cx46), connexin-50 (Cx50), and heterotypic Cx46/50 gap junction channels based on high-resolution open-state structures, enabling characterization of both ion permeation and long-timescale channel dynamics. Simulations reveal a multi-ion, multi-pathway permeation mechanism governed by isoform-specific energetic barriers and transient ion coordination sites that shape conductance and selectivity. In heterotypic Cx46/50 channels, asymmetric energetic landscapes establish a mechanistic basis for rectification. Unexpectedly, the microsecond-timescale simulations further revealed dynamic interactions between the intracellular loop (ICL) region and N-terminal domain (NT) that transiently constrict the pore and attenuate ionic currents. These findings suggest that the open-state comprises an ensemble of rapidly interconverting conductance microstates rather than a single static conformation, providing structural information of potential mechanistic importance beyond what has been learned from cryo-EM studies. Together, our results provide a mechanistic basis for ion permeation and current modulation in gap junction channels and highlight the importance of long-timescale protein dynamics in shaping intercellular communication.

## INTRODUCTION

Our mechanistic understanding of ion channel conductance, selectivity, and gating has been shaped largely by studies of narrow-pore channels, including voltage- and ligand-gated ion channels^1–3^. These systems conduct ions through highly constrained permeation pathways that range from highly selective (e.g., KcsA^4^) to relatively non-selective (e.g., TRPV1^5^). More recently, structural insights have emerged for the family of large-pore ion channels, including connexins^6–9^, innexins^10^, pannexins^11^, calcium homeostasis modulator proteins (CALHMs)^12^, and volume-regulated anion channels (VRACs)^13^. Unlike classical ion channels, these proteins support the passage of ions and larger metabolites through wide, solvent-accessible pores^14^. However, the molecular principles governing permeation and selectivity in large-pore channels remain poorly understood.

Among the members of the large-pore ion channel family, the connexins are the most functionally and structurally characterized, yet the molecular basis of their permeation, selectivity, and gating remains poorly resolved. Connexins form gap junction channels that directly couple adjacent cells, enabling rapid and direct intercellular exchange of ions, metabolites, and signaling molecules. These channels support synchronized electrical activity in excitable tissues and maintain homeostasis throughout the body^15,16^. Consequently, connexin dysfunction is associated with numerous human diseases, including neuropathies, cardiac arrhythmias, deafness, and blindness^17,18^. Gap junction channels are assembled by docking two hexameric hemichannels from neighboring cells, forming a dodecameric intercellular channel with a ∼1.2 nm wide pore^19,20^. Despite their large-pore architecture, different connexin isoforms display substantial variability in conductance, ion selectivity, and permeability to small molecules^21,22^.

A defining feature of connexins is that the N-terminal domain (NT), which shapes channel properties and participates in voltage-dependent and chemical gating, is integrated directly into the ion-permeation pathway^19,23–31^. Unlike classical voltage-gated channels, which employ specialized voltage-sensing domains distinct from the pore, connexins use the NT itself to couple changes in membrane voltage with ion permeation^6,7,28,32^. The intracellular loop (ICL) has also been implicated in channel regulation^33,34^, although its mechanistic role remains poorly understood due to intrinsic disorder. Functional studies have further shown that channel properties vary substantially among connexin isoforms and are influenced by heteromeric and heterotypic assembly, giving rise to diverse permeation behaviors, including rectification^35–37^. Yet, the molecular mechanisms underlying these properties remain incompletely resolved.

The crystal structure of connexin-26 (Cx26) provided the first atomic model of a gap junction channel in a putative open state^32^ and served as a foundation for subsequent structural and computational studies. However, later molecular dynamics (MD) simulations suggested that this structure may not represent a stable conductive state due to instability of the NT domain^7,32,38^. More recent cryo-EM structures of lens connexins Cx46 and Cx50 resolved stable open-state conformations of the NT in native heteromeric and homomeric channels^6,7^. The functional relevance of these structures has been supported by MD simulations and structure-guided mutagenesis studies^7,28^, which demonstrate the NT as an important determinant of conductance and selectivity.

Nevertheless, fundamental questions remain regarding how ions traverse these wide-pore channels, how channel asymmetry gives rise to selectivity and rectification, and how long-timescale protein dynamics shape the conductive state.

To address these questions, we focused on the closely related lens connexins Cx46 and Cx50, which share a high degree of sequence and structural similarity yet exhibit distinct conductance, selectivity, and rectification properties. This combination provides a powerful system for dissecting how subtle sequence differences shape ion permeation and channel function. We therefore performed ∼67 µs of all-atom MD simulations of homomeric Cx46 and Cx50 channels, as well as heterotypic Cx46/50 channels, under equilibrium and applied transjunctional voltage (*V*_j_) conditions. These simulations reveal a multi-ion, multi-pathway permeation mechanism shaped by isoform-specific energetic landscapes and transient ion coordination sites that govern conductance, selectivity, and rectification. In addition, long-timescale simulations uncovered dynamic interactions between regions of the ICL and NT domain that suggest a previously unrecognized mechanism for modulating open-state conductance. Together, these findings provide new insight into the molecular basis of ion transport through gap junction channels and establish general principles underlying permeation and regulation in large-pore ion channels more broadly.

## RESULTS

### K^+^ conductance involves multiple transient coordination sites

To investigate the molecular basis of *V*_j_-driven ion permeation in homomeric Cx46 and Cx50 gap junction channels, all-atom MD simulations were performed using high-resolution cryo-EM structures^6^. Each dodecameric channel was embedded in dual POPC membranes, solvated with 150 mM intracellular KCl and extracellular NaCl, and subjected to *V*_j_ ranging from 0–200 mV (**Fig. 1A**; **Fig. S1**; **Table S1**). Shorter-timescale simulations (250 ns; *n* = 4 per condition) were used to characterize conductance and permeation properties, whereas longer-timescale simulations (∼2 µs; *n* = 3 per condition) were used to explore extended dynamics and rare events. All systems reached steady-state behavior within ∼50 ns and maintained stable open-state NT conformations throughout the simulations^7^ (**Figs. S2, S3**).

**Figure 1.**
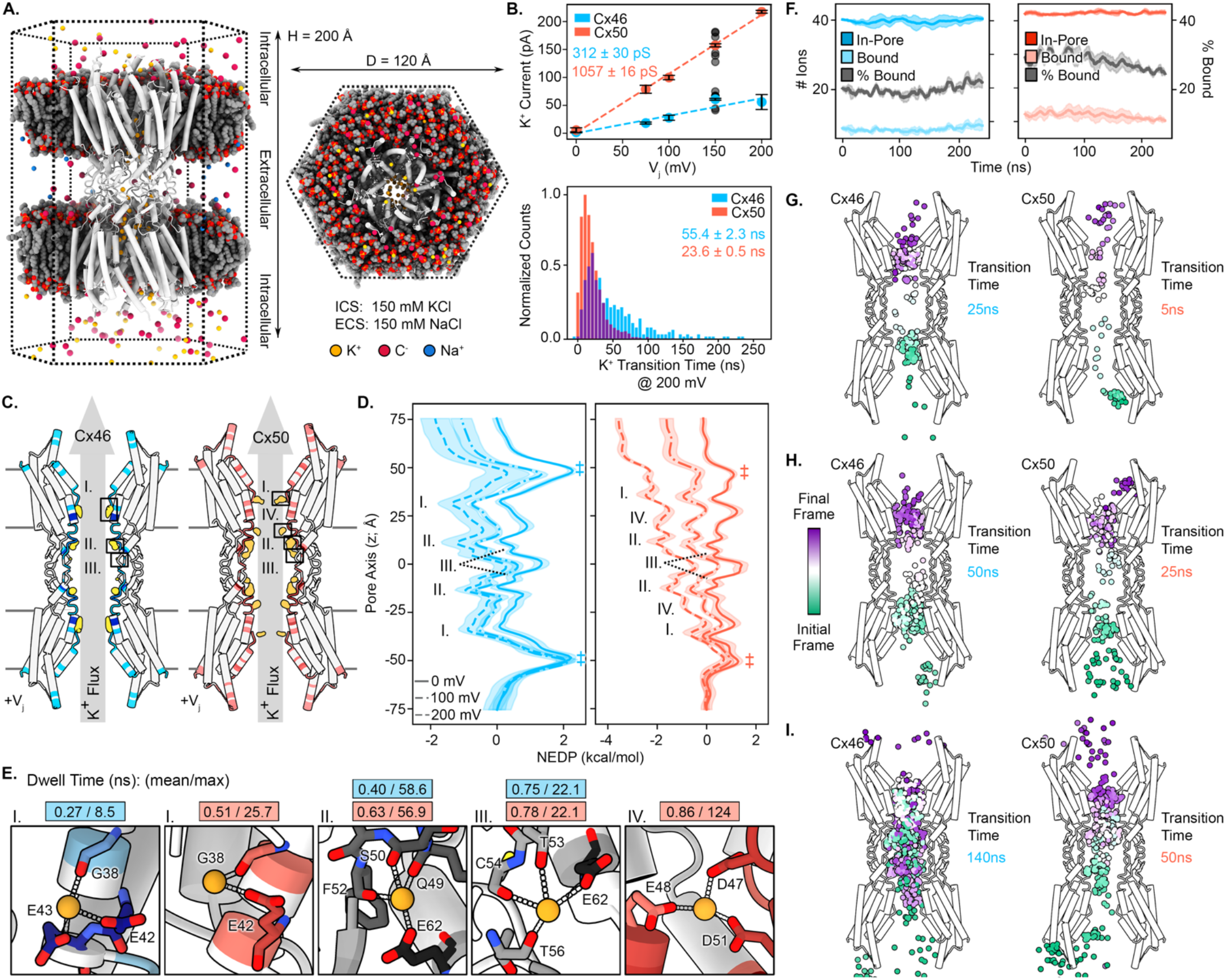
Vj-driven K+ permeation in Cx46 and Cx50 gap junctions. **A**) Hexagonal simulation box depicting Cx46 (white) embedded in dual POPC bilayers (grey and red). Ions are colored as K^+^ (yellow), Cl^−^ (red) and Na^+^ (blue); explicit water molecules omitted for clarity. **B**) Current-Voltage (*I/V*_j_) relationships for Cx46 (blue) and Cx50 (red) at various *V*_j_ potentials (top). Error bars represent s.e.m.; black circles at *V*_j_ = 150 mV indicate 250 ns blocks from long-timescale simulations. Distribution of K^+^ transition-times (ns) for Cx46 (blue) and Cx50 (red) at *V*_j_ = 200 mV (bottom). **C**) Cross-sectional views of Cx46 (left) and Cx50 (right), with time-averaged K^+^ densities (yellow) highlighting interaction sites: I-III for Cx46 and I-IV for Cx50. Proteins are colored by K^+^ contact frequency. **D**) Non-equilibrium driving potentials (NEDPs) for K^+^ permeation through Cx46 (left, blue) and Cx50 (right, red), depicting the energetic landscape under varying *V*_j_. Energy minima correspond to K^+^ interaction sites, labeled. Entrance and exit barriers annotated (‡). **E**) Close-up views of K^+^ interaction sites in Cx46 (blue) and Cx50 (red), with coordinating residues are labeled. Binding dwell times (ns) at *V*_j_ = 200 mV are displayed (mean/max values). A single representative structure is shown for interaction sites II and III, which are conserved between Cx46 and Cx50. **F**) Number of ions in the pore (dark traces) and bound to the protein (light traces) as a function of time for Cx46 (left, blue) and Cx50 (right, red) at *V*_j_ = 200 mV. The percentage of bound ions is shown (gray traces, right axis). Data represent mean ± s.e.m. (n=4). **G-I**) Snapshots from MD trajectories illustrating fast (G), medium (H), and slow (I) K^+^ permeation pathways through Cx46 (left) and Cx50 (right). Ion coordinates are colored by trajectory frame. Transition times correspond to the permeation events are annotated.

Both isoforms exhibited linear *I/V* relationships (r^2^_Cx46_ = 0.89, r^2^_Cx50_ = 0.99), with slope junctional K⁺ conductance values (*γ*_j_) of 312 ± 31 pS and 1057 ± 16 pS, respectively. Consistent with these conductance differences, K^+^ transition times – first-passage times excluding the time to enter the pore – were significantly slower in Cx46 than Cx50 (52.4 ± 20 ns versus 24.2 ± 8 ns at *V*_j_ = 200 mV; p < 0.001) (**Fig. 1B**). These findings are consistent with the lower conductance of Cx46 relative to Cx50 observed experimentally^37,39^. Notably, however, simulated conductance values exceeded reported single-channel measurements by ∼2–5 times, a discrepancy explored in later sections.

To identify the molecular determinants of permeation, time-averaged K⁺ densities and non-equilibrium driving potentials (NEDPs) were calculated from *V_j_* = 200 mV simulations (**Fig. 1C,D**). K⁺ ions occupied a series of transient coordination sites distributed along electronegative regions of the pore lumen, forming a multi-site route of permeation rather than a single continuous pathway. Four major interaction sites (I–IV) were identified within the NT vestibule, TM1, and EC1 regions (**Figs. 1C–E; S4–S5**).

Sites I–III were conserved between Cx46 and Cx50. Site I was coordinated primarily by E42 and G38, with additional fluctuating contributions from neighboring residues including D3, as well as E43 in Cx46 (F43 in Cx50) (**Fig. 1E**). Site II exhibited the highest occupancy and formed a prominent energetic minimum within the permeation pathway. This site was coordinated by E62 together with backbone carbonyls from the EC1 loop. Site III was positioned immediately adjacent to Site II and formed a contiguous cation-interaction region within EC1. Notably, Site II overlaps with a recently identified Ca^2+^ binding site in Cx46/50^30^, suggesting that the same structural elements may support both monovalent permeation and divalent ion regulation.

A fourth interaction site (Site IV) was observed predominantly in Cx50 and was coordinated by E47, D48, and D51 (**Fig. 1C–E**). Although these residues are conserved in Cx46, alternative sidechain conformations reduced site occupancy and dwell times relative to Cx50 (**Fig. S2; S4**). Several residues contributing to Sites I-IV, including D3, E42, E43, Q49, D51, and E62, have previously been implicated in connexin conductance by mutagenesis studies^7,21,40,41^, further supporting the functional relevance of these coordination sites. Consistent with this interpretation, time-averaged ion densities aligned closely with energetic minima in the NEDP profiles (**Fig. 1D**), indicating that these regions function as transient energy wells that facilitate K^+^ permeation.

The most prominent isoform-specific difference occurred near the NT vestibule, where residue 9 shapes the entrance and exit barriers encountered by permeating cations. In Cx46, the positively charged R9 imposed significantly larger energetic barriers than the equivalent uncharged N9 residue in Cx50 (**Fig. 1D**; entrance/exit barriers of 2.1 ± 0.5 and 1.6 ± 0.8 kcal/mol in Cx46 versus 0.8 ± 0.2 and 0.1 ± 0.2 kcal/mol in Cx50; p < 0.05, p < 0.01), accounting for much of the reduced K^+^ conductance observed in Cx46. Both channels also exhibited a smaller central barrier near z = 0 Å, consistent with a region of lower coulombic potential and previous equilibrium-state models (**Fig. S5**)^7,28^. Increasing *V*_j_ reduced the exit barrier in both channels, with a larger effect in Cx46 (0.5 kcal/mol vs. 0.2 kcal/mol in Cx50), likely reflecting partial compensation of the electrostatic repulsion generated by R9. Thus, isoform-specific differences in transient coordination sites and vestibular electrostatics shape the permeation landscape and account for the distinct K^+^ conductance properties of Cx46 and Cx50.

### Ion permeation occurs via a multi-ion, multi-path mechanism

In contrast to classical ion channels that support single-file permeation through narrow, highly selective filters, Cx46 and Cx50 possess large, solvent-filled pores that permit multiple ions to permeate simultaneously via diverse pathways. These wide pores allow ions to dynamically sample a network of transient interactions with the pore-lining residues (**Fig. 1F–I**).

Across simulations, both Cx46 and Cx50 maintained an average of ∼40 K^+^ ions within the pore under steady-state conditions, indicating substantial ion occupancy under applied *V*_j_ potentials. Based on identified binding sites across all subunits, up to 36 and 48 potential K^+^ coordination sites were mapped for Cx46 and Cx50, respectively. However, only a subset of these sites were occupied at any given time, with approximately 20% in Cx46 and 28% in Cx50, regardless of *V*_j_ magnitude (**Fig. 1F**). The higher binding site occupancy in Cx50 reflects a higher apparent affinity for K^+^, consistent with the enhanced ion occupancy and longer dwell times observed at its binding sites (**Fig. S4**).

K^+^ permeation displayed a wide range of transition times (**Fig. 1B***, bottom*), with some ions taking over 100 ns to traverse the channel, while others completed translocation in as little as 5 ns. These broad distributions with long, non-uniform tails reflect diverse permeation trajectories shaped by transient binding, energetic barriers, and inter-ion interactions. Typically, ions encountered energetic barriers at the channel center or cytoplasmic exit, including electrostatic interactions with negatively charged residues hinder forward progress (**Fig. 1G–I; Movie S1**).

Interestingly, the fastest K^+^ transitions, particularly in Cx50, occurred with minimal protein contact, as ions effectively “surfed” through the channel in the presence of other bound ions (**Fig. 1G; Movie S2**). This effect appears to arise from electrostatic shielding, where resident K^+^ ions transiently occupy nearby binding sites, diminishing local electrostatic attractions and preventing subsequent ions from becoming energetically trapped. These results support a multi-ion, multi-pathway mechanism in which shallower entrance and exit barriers, together with greater occupancy-dependent electrostatic shielding, allow K⁺ ions to traverse Cx50 more efficiently than Cx46.

### Monovalent cations follow a common permeation mechanism

Cx46 and Cx50 exhibit monovalent cation selectivity in electrophysiology experiments consistent with the classical permeability sequence Cs+ > K+ > Na+, which parallels their relative diffusion coefficients in bulk solution^19,42^. Simulations performed at *V*_j_ = 200 mV under different ionic conditions (150 mM CsCl, KCl, and NaCl) reproduced this selectivity sequence in both isoforms (**Fig. S3,S6; Table S1**). While conductance varied among ions, Cs^+^, K^+^, and Na^+^ occupied similar coordination sites and traversed comparable energetic landscapes within the pore.

The reduced Na^+^ current relative to K^+^ is broadly consistent with its lower diffusion coefficient in bulk solution (D_Na_/D_K_ = 0.68)^43^. In contrast, Cs^+^ conductance exceeded predictions based on bulk diffusion alone (D_Cs_/D_K_ = 1.05), suggesting a more favorable permeation pathway. Consistent with this interpretation, Cs^+^ experienced a shallower entrance barrier than K^+^, likely reflecting weaker interactions with negatively charged residues near the cytoplasmic vestibule, including those within the ICL region (**Fig. S6**).

Despite differences in conductance, Cs^+^ and Na^+^ followed permeation mechanisms similar to K^+^, engaging comparable coordination sites and energy landscapes along the channel interior (**Fig. S6**). Given this overall similarity, KCl was used as the standard condition for comparative analyses in the remainder of the study.

### Cl^−^ follows a distinct permeation mechanism

Cx46 and Cx50 exhibit modest cation selectivity^21,42^, indicating that both cationic and anionic flux contribute to overall channel conductance. However, unlike for K^+^, Cl⁻ permeation events were rare in short-timescale simulations (∼250 ns), necessitating extended sampling. Long-timescale simulations (∼2 µs) performed at *V*_j_ = 150 mV captured sufficient permeation events to characterize the Cl⁻ pathway in both channels (**Fig. 2**).

**Figure 2.**
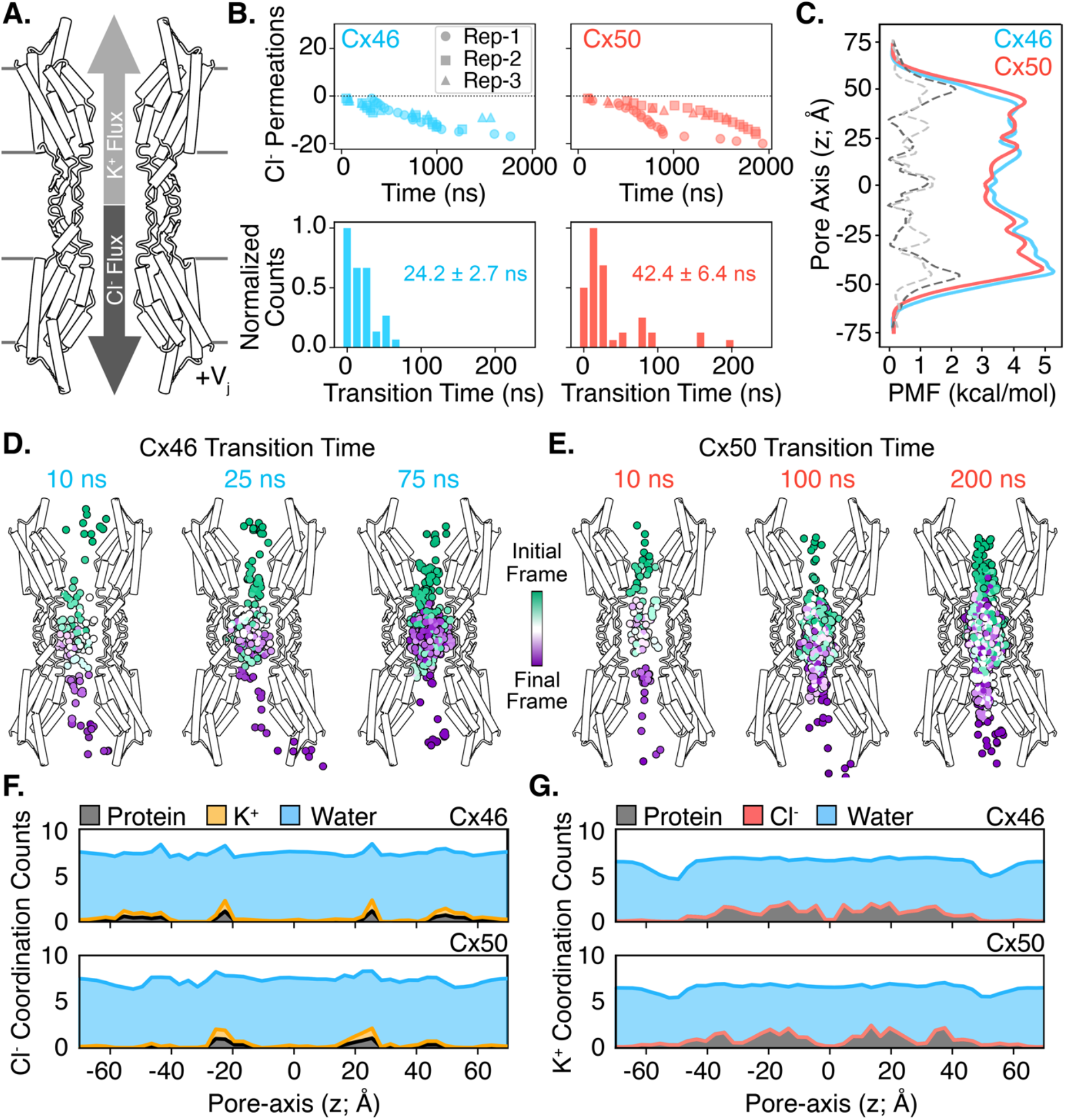
Vj-driven Cl– permeation in Cx46 and Cx50 gap junctions from long timescale simulations. **A**) Schematic cross-section of a gap junction (Cx46 or Cx50) illustrating simultaneous counter-currents of K^+^ and Cl^−^. **B**) Cl^−^ flux traces for Cx46 (left) and Cx50 (right) from long-timescale simulations at *V*_j_ = 150 mV (top). Corresponding Cl^−^ transition-time distributions, with mean *±* s.e.m. values displayed (bottom). **C**) Potentials of mean force (PMF) for Cl^−^ ions calculated from a single combined Markov State Model (MSM) for Cx46 (blue) and Cx50 (red). Due to limited Cl^−^ sampling, Independent MSMs were not generated for each replicate. Mean PMFs for K^+^ at *V*_j_ = 150 mV are shown for comparison (Cx46, light gray dashed; Cx50, dark gray dashed). **D,E**) Representative Cl^−^ permeation pathways illustrating fast, medium, and slow transitions for Cx46 (D) and Cx50 (E), with respective transition times indicated. Ion coordinates are colored by trajectory frame. **F,G**) Cumulative counts of Cl^−^ (F) and K^+^ (G) coordination with protein (grey), water (blue), K^+^ (yellow) and Cl^−^ (red), as a function of the pore axis in Cx46 (upper) and Cx50 (lower), derived from trajectories at *V*_j_ = 150 mV (n = 3).

Cl⁻ ions permeated in the direction opposite to K^+^, generating a *V*_j_-driven counter-current (**Fig. 2A**; **Movie S3**). The first permeation events exhibited a substantial lag (∼50–100 ns) after the start of simulations, approximately tenfold longer than observed for K^+^, indicating the presence of a significant entrance barrier (**Fig. 2A,B**; **Fig. S10**). Once inside the pore, Cl⁻ traversed the channel on a timescale comparable to K+, although with a reversed isoform dependence (19.8 ± 11 ns for Cx46 and 33.7 ± 25 ns for Cx50). This reversal mirrors the relative K+/Cl⁻ selectivity of the two channels and suggests that entry into the pore, rather than transit through its interior, represents the principal kinetic bottleneck for anion permeation.

This behavior was reflected in the energetic landscape. Potential of mean force (PMF) profiles revealed entrance barriers of ∼4–5 kcal/mol for Cl⁻, substantially larger than those observed for K^+^ (**Fig. 2C; Fig. S10**). These elevated barriers likely arise from electrostatic repulsion by negatively charged residues lining the NT vestibule and EC1 region. Consistent with this interpretation, removal of N-terminal acetylation modestly decreased the Cl^−^ entrance barrier while increasing the barrier for K^+^ permeation (**Fig. S10**), further supporting a role for NT electrostatics in shaping connexin selectivity. Following entry, Cl^−^ occupied a broad shallow energy well within the central cavity before overcoming cumulative barriers of ∼2–3 kcal/mol, resulting in periods of transient stalling that further limited anion flux.

Trajectory analysis revealed a permeation mechanism fundamentally distinct from that of K^+^. Whereas K^+^ permeated through a distributed network of transient coordination sites, Cl^−^ exhibited few stabilizing interactions with pore-lining residues and remained predominantly water-coordinated throughout transit (**Fig. 2D–G**). Contacts with the protein or neighboring ions were infrequent and transient, indicating that Cl^−^ permeation is governed primarily by electrostatic exclusion rather than favorable protein-ion interactions. The modest cation selectivity of Cx46 and Cx50 therefore appears to arise primarily from energetic exclusion of Cl⁻ at the channel entrance rather than strong discrimination among monovalent cations. Cations and anions thus traverse the same pore through fundamentally different mechanisms.

### Isoform-specific energetic barriers underlie heterotypic Cx46/50 rectification

Rectification refers to directional bias in ion conduction that depends on the applied voltage potential. While homomeric gap junction channels typically exhibit symmetric conductance due to their internal symmetry, heterotypic channels formed from distinct connexin hemi-channels can display pronounced rectification that can direct the flow of ionic and chemical communication across tissues^35,36,44,45^. To investigate the molecular basis of this behavior, a heterotypic Cx46/50 channel was constructed and simulated across a *V*_j_ series (±75 to ±200 mV), with polarity defined relative to the Cx46 side (**Fig. 3A,B; Table S1**).

**Figure 3.**
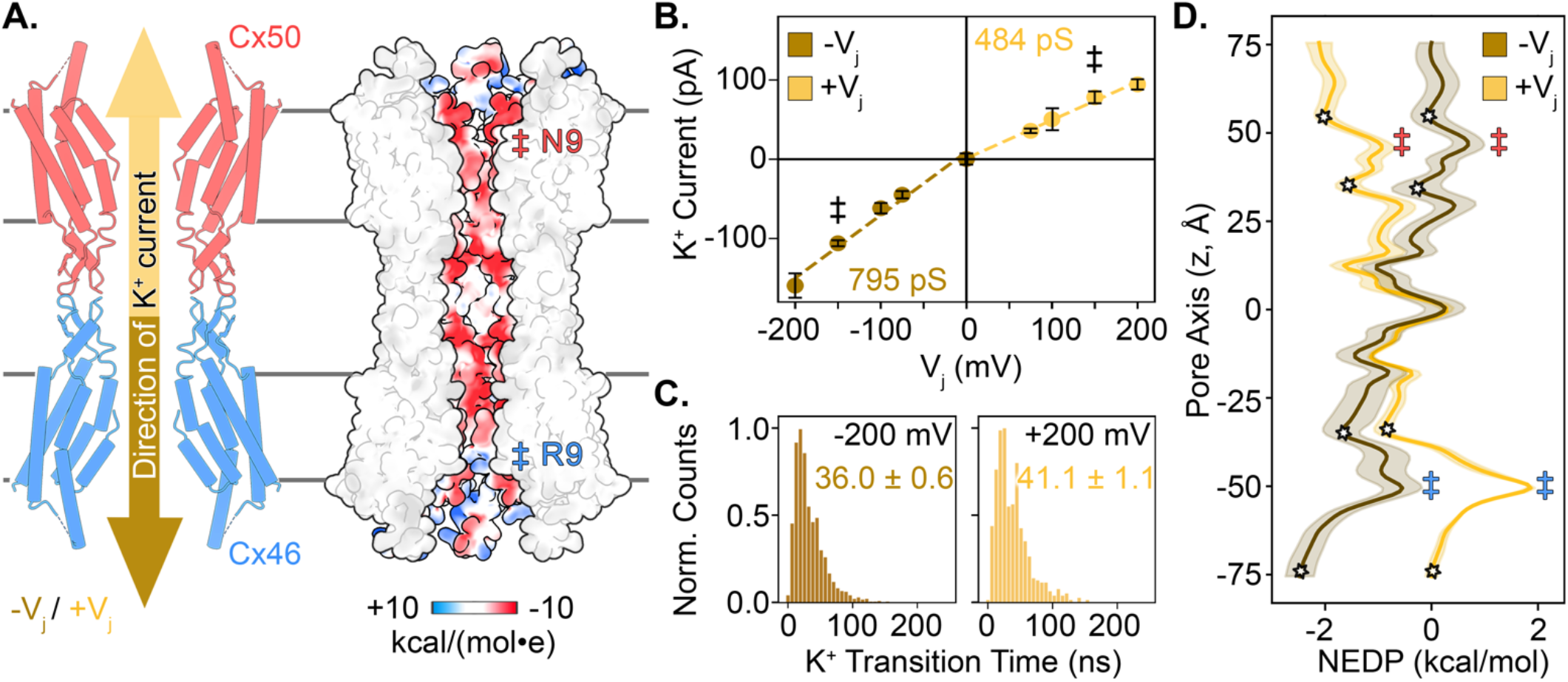
Rectifying K+ currents in Cx46/50 heterotypic gap junctions. **A**) Cross-sectional diagram of the heterotypic Cx46/50 gap junction, showing the Cx46 hemichannel (blue) and Cx50 hemichannel (red), along with K^+^ flux directions under negative (-*V*_j_, brown) and positive (+*V*_j_, yellow) transjunctional voltage (left). Coulombic surface potential of the heterotypic Cx46/50 channel, shown in split-view to highlight the electrostatic properties of the permeation pathway (right). Coulomb potential displayed as positive (blue), neutral (white), and negative (red). Amino acid positions of R9 (Cx46) and N9 (Cx50) indicated (‡). **B**) K^+^ current-voltage (*I/V*_j_) relationship for heterotypic Cx46/50 over a *V*_j_ range of -200 mV to +200 mV. Simulations with ± 150 mV come from long timescale Anton2 trajectories (‡). Data represent mean ± s.e.m from short-timescale (n=4) and long-timescale (n=3) simulations. **C**) K^+^ transition time distributions at *V*_j_ = -200mV (left) and +200 mV (right), with mean ± s.e.m. values displayed. **D**) Non-equilibrium driving potentials (NEDPs) for heterotypic Cx46/50 channels at *V*_j_ = -200 mV (brown) and +200 mV (yellow), presented as mean ± s.e.m. (n=3 replicates). Entrance and exit barriers are annotated (‡), with energy minima used for barrier calculations indicated (stars).

The heterotypic channel exhibited clear rectification, with larger K^+^ currents at negative *V*_j_ (corresponding to ion entry through Cx50) than at positive *V*_j_ (**Fig. 3B**). Although absolute conductance values exceeded experimental measurements, as observed with the homomeric simulations, the degree of rectification of ∼1.6 (conductance ratio at ± 200 mV) was similar to that observed experimentally of ∼1.5^37^. K^+^ transition times further reflected this asymmetry, with significantly slower permeation at +200 mV (41.1 ± 1.1 ns) than at −200 mV (36.0 ± 0.6 ns; p < 0.001) (**Fig. 3C**).

The basis of rectification was revealed by the NEDPs derived from the same trajectories (**Fig. 3D**). At positive *V*_j_, K^+^ ions entering through the Cx46 hemichannel encountered a substantially larger entrance barrier than ions entering through Cx50 at negative *V*_j_. This asymmetry originates primarily from residue 9 within the NT domain, where a positively charged arginine in Cx46 (R9) and a neutral asparagine in Cx50 (N9) establish distinct entrance barriers in the corresponding homomeric channels. When combined within a heterotypic channel, these isoform-specific differences create an asymmetric energetic landscape for permeation. Thus, rectification emerges directly from the unequal energetic costs associated with ion entry through the two hemichannels.

The applied *V*_j_ further amplified this asymmetry by selectively modulating the Cx46 exit barrier. At z = −50 Å, the exit barrier decreased from ∼1.6 kcal/mol in the absence of an applied voltage to ∼0.8 kcal/mol at *V*_j_ = −200 mV (**Fig. 3D**), likely reflecting partial compensation of the electrostatic repulsion imposed by R9. Similar voltage-dependent effects were observed in homomeric Cx46, indicating that rectification arises not only from intrinsic isoform-specific differences in permeation barriers, but also from their differential response to the applied electric field. Rectification in Cx46/50 channels therefore emerges from asymmetric energetic landscapes established by isoform-specific electrostatics at the NT vestibule, while differential voltage-dependent modulation of these barriers further amplifies directional ion flux.

### Modulation of gap junctional currents by a dynamic ICL-NT interactions

MD simulations reproduced the relative conductance, selectivity, and rectification properties of Cx46 and Cx50, but the shorter-timescale simulations (∼250 ns) consistently yielded higher conductance values than measured experimentally. Although channels remained globally stable and currents reached an apparent steady state over these simulations, extending the simulations to ∼2 μs unexpectedly revealed a progressive attenuation of ionic currents in Cx46, Cx50, and heterotypic Cx46/50 channels (**Fig. 4A,B; Figs. S7–S8**).

**Figure 4.**
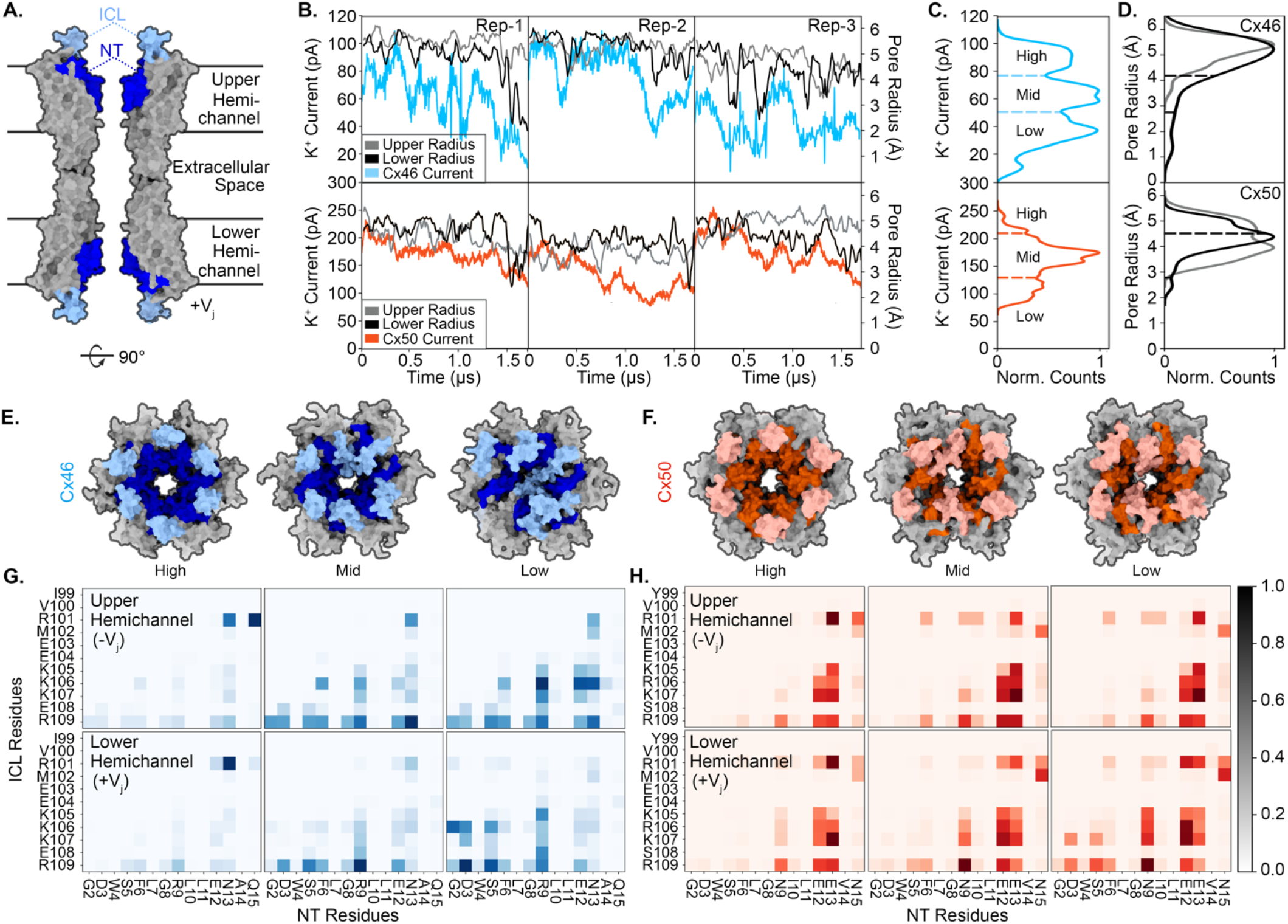
Attenuation of ion currents by dynamic ICL–NT interactions in Cx46 and Cx50. **A**) Cross-sectional view of Cx46 in surface representation, highlighting the NT domain (dark blue) and modeled region of the ICL (light blue). Upper and Lower hemichannels are labeled for reference to *V*_j_ polarity. **B**) Traces of K^+^ current (left axis) and minimum pore radius (right axis) over time for independent Anton2 replicates (n = 3; Rep-1 to Rep-3). Window-averaged K^+^ currents (100 ns windows) are shown in color (Cx46, blue; Cx50, red), with the Upper and Lower pore radii overlaid in gray and black, respectively. **C**) Distribution of K^+^ currents over time, clustered as ‘High, ‘Mid’, and ‘Low’ conductance states (dashed lines). **D**) Distributions of the Upper and Lower pore radii, clustered corresponding to the conductance-states in Panel C (dashed lines). **E,F**) Views of the channel pore from the mean structure for each conductance state in Cx46 (E) and Cx50 (F). **G,H**) ICL–NT contact maps for Upper (top) and Lower (bottom) hemichannels in Cx46 (G) and Cx50 (H), corresponding to each conductance state. Color intensity represents the normalized interaction frequency.

Current attenuation correlated closely with narrowing of the pore radius (r_min_), defined as the minimum radius within each respective hemichannel, which decreased from ∼5 Å in the high conductance open-state to ∼1.25– 2.5 Å in the most attenuated states (**Fig. 4B**). Clustering of current amplitudes identified high-, mid-, and low-conductance states that closely tracked corresponding r_min_ distributions (**Fig. 4C,D**). In the lowest conductance states, pore constriction exceeds the hydrated radius of K^+^ (3.3 Å)^46^, consistent with substantial reduction of ionic flux.

To determine the structural basis of this attenuation, representative structures were analyzed for each conductance state. These revealed progressive inward movement of a modeled segment of the intracellular loop (ICL; residues 99–109) extending from TM2 (**Fig. 4E,F**). While the full-length ICL was not resolved in available cryo-EM structures and therefore could not be modeled, the resolved proximal segment exhibited striking state-dependent behavior. In high-conductance states, the ICL retained an α-helical conformation similar to the open-state cryo-EM structures^6,7^. In mid- and low-conductance states, the ICL became increasingly disordered and extended toward the pore lumen, forming transient interactions with the NT domain. In Cx46, extensive ICL-NT interactions produced near-complete occlusion of the cytoplasmic vestibule, whereas similar but reversible interactions were observed in Cx50 and heterotypic channels (**Fig. S8; Movie S4**).

Residue-contact analysis further demonstrated progressive engagement of the NT by distal ICL residues during conductance attenuation (**Fig. 4G,H**). Whereas high-conductance states were characterized by limited contacts between the proximal ICL and distal NT, low-conductance states displayed extensive interactions involving residues near the NT constriction site (positions 2–9). The deepest penetrating interaction involved R109 and D3, which correlated with collapse of the cytoplasmic vestibule and substantial current reduction.

To assess voltage dependence, additional long-timescale simulations were performed in the absence of applied *V*_j_ (**Fig. S9**; **Table S1**). Cx46 maintained a stable open pore in all replicates, with r_min_ values of ∼5 Å, whereas spontaneous ICL-NT interactions and partial pore closure were observed in one Cx50 replicate and on the Cx50 side of heterotypic channels. These observations indicate that ICL-NT interactions are not strictly voltage-dependent, although the applied electric field may stabilize or enhance their formation.

These results indicate that the open-state of connexin channels comprises an ensemble of rapidly interconverting conductance microstates rather than a single stable conformation. Transient ICL-NT interactions can act as a dynamic attenuator of ionic current by modulating pore accessibility on the microsecond timescale. This behavior may contribute to the lower conductance values measured experimentally and reveals a previously unrecognized mechanism through which long-timescale protein dynamics regulate gap junction channel function.

## DISCUSSION

Using long-timescale all-atom simulations, we identify the energetic and dynamic mechanisms that govern ion permeation, selectivity, rectification, and conductance modulation in Cx46 and Cx50 gap junction channels. The results support a multi-ion permeation mechanism shaped by isoform-specific electrostatics and transient ion coordination sites and suggest a previously unrecognized role for ICL dynamics in modulating open-state conductance. Together, these findings demonstrate that voltage-driven ion permeation and long-timescale protein dynamics contribute fundamentally to gap junction function in ways that are not captured by static structural snapshots.

### Multi-Ion, multi-pathway permeation in a large-pore channel

Unlike classical ion channels that support single-file permeation through narrow selectivity filters, Cx46 and Cx50 contain wide, solvent-accessible pores capable of accommodating dozens of ions simultaneously. Rather than traversing a single defined pathway, ions permeate through a series of transient coordination sites and electrostatic barriers that collectively shape the energetic landscape of the channel (**Fig. 5A**). These findings suggest that permeation through connexins is governed by fundamentally different principles than those established for classical narrow-pore ion channels.

**Figure 5.**
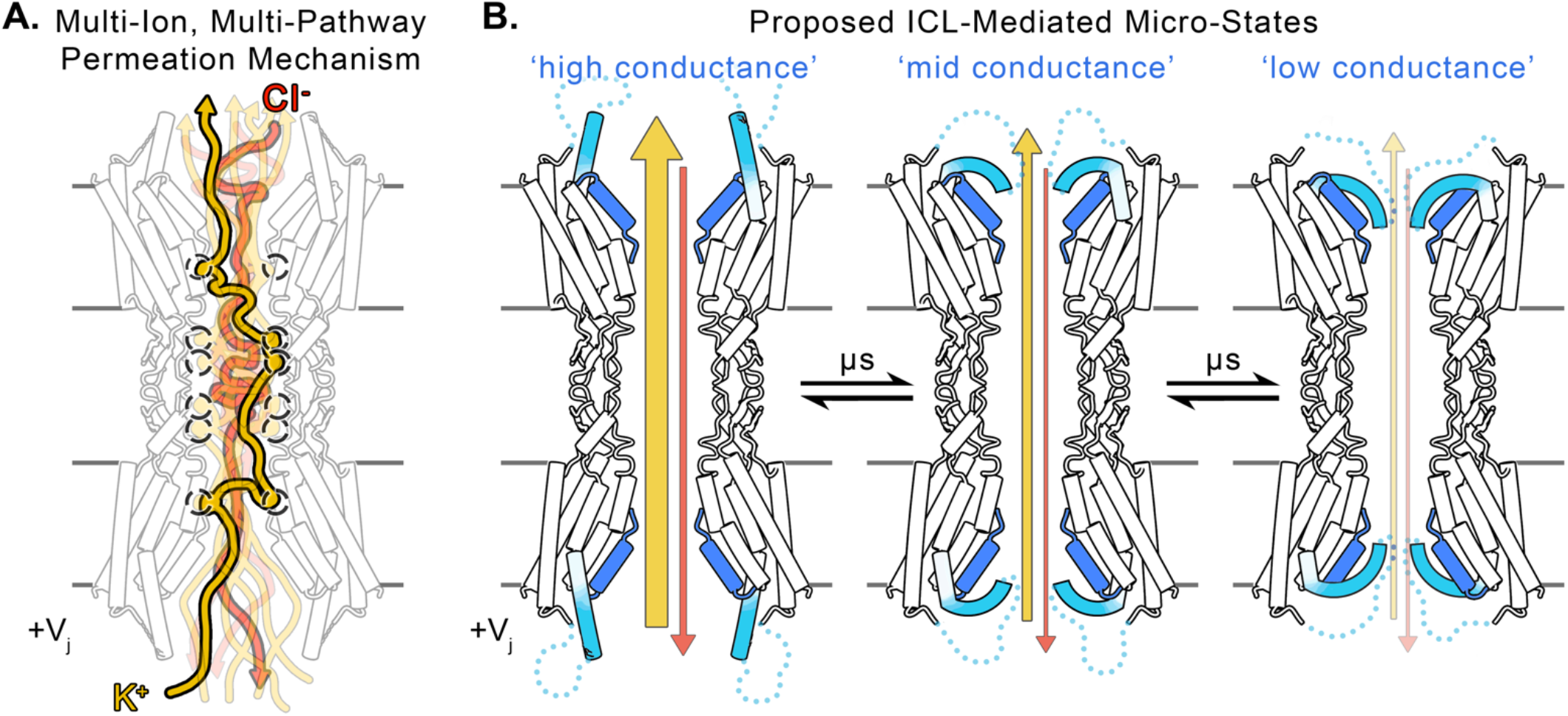
Overview of multi-ion, multi-pathway permeation mechanism and ICL-mediated current attenuation. **A**) Schematic representation of the multi-ion, multi-pathway permeation mechanism in Cx46 and Cx50, as observed in long-timescale MD simulation. The large-pore gap junction channel is occupied by multiple monovalent cations (e.g., K^+^, yellow) and anions (e.g., Cl^−^, red), which follow distinct permeation pathways driven by the transjunctional voltage potential (*V*_j_). Due to the electronegative coulombic potential inside the pore, Cx46 and Cx50 preferentially permeate monovalent cations over anions. Weak cation interaction sites (black dashed circles) are transiently occupied during K^+^ permeation. More efficient permeation occurs due to site occupancy, while slower permeation events involve interactions with multiple binding sites. Cl^−^ counter-currents flow in the opposite direction of cations, with permeation largely dictated by the large entrance barrier, and a smaller exit barrier leaving central vestibule, following pathways of minimal interaction with the negatively charged pore lumen. **B**) Long-timescale MD simulations revealed a dynamic ICL-NT interaction network that attenuates ion conductance. Simulated ion conductance for Cx46 and Cx50 was significantly higher than expected based on the fully open-state conformation of the starting cryo-EM structures. However, the proximal ICL region (light blue) exhibited transient interactions with the NT domain (dark blue), leading to graded conductance states: high conductance (minimal ICL-NT interactions), mid conductance (moderate ICL-NT interactions), and low conductance (substantial ICL-NT interactions). The complete ICL was not included in the MD simulations due to its absence in the cryo-EM structures, where it remains unresolved as an intrinsically disordered domain (light blue dotted lines). The observed ICL-NT fluctuations occur on a microsecond timescale, suggesting the ensemble of these ‘micro-states’ may contribute to an overall attenuation of ionic currents, aligning more closely with conductance values of the open-state measured on the millisecond timescale in single-channel electrophysiological recordings.

The higher K^+^ conductance of Cx50 relative to Cx46 appears to arise not from increased pore capacity, but from a more permissive energetic landscape. Reduced entrance barriers and enhanced occupancy of transient coordination sites promote productive ion transit while limiting energetic trapping within the pore. Because these sites are frequently occupied under multi-ion conditions, permeating K^+^ ions often bypass otherwise favorable interaction sites, effectively "surfing" over transient energetic minima rather than becoming sequentially trapped at each site. Notably, the most frequently occupied coordination site overlaps with a recently identified Ca^2+^ binding site in Cx46/50^30^, suggesting that ion coordination sites within the permeation pathway may serve dual roles in ion conduction and regulation. The ability of both monovalent and divalent cations to occupy these sites raises the possibility that they represent conserved structural elements through which connexin channels tune permeation and gating behavior.

Cl^−^ permeation followed a fundamentally different mechanism. Relative to K^+^, Cl^−^ encountered substantially larger entrance barriers, exhibited few stabilizing interactions with pore-lining residues, and remained predominantly water coordinated throughout transit. These observations suggest that the modest cation selectivity of Cx46 and Cx50 arises primarily from energetic exclusion of Cl^−^ rather than preferential stabilization of permeating cations. Consistent with previous studies, the state of N-terminal acetylation further influenced channel conductance and selectivity, supporting a broader role for post-translational modification of the NT domain in tuning connexin channel properties^7,47^. Together, these observations support a model in which permeation through connexin channels is governed by a multi-ion, multi-pathway mechanism distinct from the single-file transport paradigms established for classical narrow-pore ion channels.

### Energetic basis of rectification in heterotypic Cx46/50 channels

Heterotypic gap junctions formed by distinct connexin hemichannels can exhibit rectification, providing a mechanism for directional intercellular signaling. Simulations of heterotypic Cx46/50 channels revealed that rectification arises from intrinsic asymmetry in the energetic landscape of the two hemichannels. Cx46 contains a positively charged R9 residue near the pore entrance that imposes a substantial energetic barrier to cation permeation, whereas the equivalent position in Cx50 is occupied by a neutral asparagine that creates a more permissive pathway. Voltage-dependent modulation of these asymmetric barriers further amplifies directional ion flux, providing a mechanistic explanation for the rectifying behavior observed experimentally in Cx46/50 channels^37,39^.

Recent work has implicated Mg^2+^ as an additional modulator of rectification through selective interactions on the Cx46 side of the channel^48^. Intriguingly, the proposed Mg^2+^ binding site overlaps with a transient cation coordination site identified in our simulations. In Cx50, this region lacks a key acidic residue present in Cx46, potentially explaining the asymmetric effects of Mg^2+^ on rectification. Together with the recently identified Ca^2+^ binding sites that overlap with the transient cation coordination sites described here^30,40^, these observations suggest that conserved ion coordination sites within the permeation pathway may serve as evolutionary hotspots through which different connexin isoforms tune conductance, selectivity, rectification, and channel gating.

### ICL-mediated attenuation of single-channel conductance

Long-timescale simulations revealed that the open-state of connexin channels is dynamically heterogeneous, with transient ICL-NT interactions attenuating ionic currents by partially occluding the cytoplasmic vestibule (**Fig. 5B**). These observations suggest that the open-state comprises an ensemble of rapidly interconverting conductance microstates rather than a single conductive conformation. Such behavior may also help reconcile the observation that conductance values predicted from relatively static structural models over shorter timescales were consistently higher than those measured experimentally.

The transitions between these microstates were mediated by electrostatic interactions between charged residues in the ICL and NT and are consistent with the pronounced conformational flexibility of the ICL observed in cryo-EM studies^6,7^. Although we do not propose that these states correspond directly to the stable sub-conductance levels observed electrophysiologically, their rapid (∼microsecond) interconversion could influence the time-averaged conductance of the open state measured on the millisecond timescale. More broadly, these findings suggest that long-timescale protein dynamics contribute directly to the conductive properties of connexin channels and that static structural models capture only one member of a broader ensemble of conductive states.

The functional importance of the ICL has remained elusive despite longstanding evidence implicating this region in connexin regulation and gating^33,34^. Our simulations provide a potential structural basis for these observations by suggesting that transient ICL-NT interactions dynamically attenuate conductance without requiring formation of a stable closed state. The observation of spontaneous pore occlusion events in simulations lacking an applied voltage further indicates that these interactions are intrinsic to the open-state ensemble, although transjunctional voltage may stabilize or enhance their formation. Together, these findings identify the ICL as an active participant in shaping open-state conductance rather than a passive structural element.

### Limitations and future directions

Several limitations should be considered when interpreting these findings. Most notably, only the proximal structurally resolved segment of the ICL was modeled, whereas the remainder of the domain is unresolved in current structures and likely highly dynamic. Additional regions of the ICL may therefore contribute to channel regulation or interactions with cytoplasmic binding partners^34^. Furthermore, while these simulations provide a detailed description of monovalent ion permeation, they do not address the mechanisms governing selective transfer of larger signaling molecules such as cAMP or IP_3_ that represent a defining feature of gap junction communication^49–51^. These larger substrates appear to exhibit saturable, non-linear kinetics that may resemble transporter-like mechanisms^49,52^, raising important questions about the conformational flexibility and energetic landscapes required for intercellular exchange of the molecules.

Future studies integrating long-timescale molecular dynamics simulations with high-resolution structural methods and single-channel electrophysiology will be essential for defining the full spectrum of connexin channel behaviors. More broadly, our findings suggest that permeation through large-pore ion channels is governed by principles distinct from those established through studies of classical narrow-pore channels. Rather than relying on highly constrained single-file transport, these channels may support ion flux through distributed energetic landscapes shaped by multi-ion occupancy, transient coordination sites, and conformational dynamics within the permeation pathway. As structural and computational studies continue to expand across connexins, pannexins, innexins, CALHM channels, and VRACs, it will be important to determine whether these features represent common principles governing ion permeation and regulation across the broader family of large-pore ion channels.

## MATERIALS AND METHODS

### MD system preparation

Dodecameric gap junction models of Cx46 (PDB ID: 7JKC) and Cx50 (PDB ID: 7JJP)^6^ were assembled using Visual Molecular Dynamics (VMD) software^53^. The heterotypic Cx46/50 model was constructed by aligning the homomeric structures and combining one hemichannel from each isoform into a single dodecamer. The N-termini were acetylated, according to mass spectrometry analysis^7^. To nullify artificial termini charges, the exposed ends of the intracellular loop (ICL) were neutralized by capping with acetylation (ACE; “n”-termini), and n-methyl amidation (NME; “c”-termini). Each channel was embedded into dual, opposing lipid bilayers composed of 1-palmitoyl-2-oleoyl-sn-*glycero*-3-phosphocholine (POPC) using CHARMM-GUI^54^, and lipid molecules overlapping with protein were removed. Systems were then solvated in a 150 × 150 × 200 Å water box using VMD’s solvate plugin^53^. The extracellular compartment contained 150 mM NaCl, while the intracellular solutions were solvated with either 150 mM KCl, NaCl, or CsCl. A summary of simulation conditions is provided in **Table S1**.

### Short-timescale simulations

Short-timescale simulations were performed locally using a hexagonal simulation box defined by the matrix:

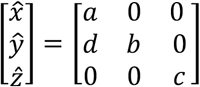

where *a* is the longest diameter 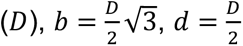 and *c* is the box height. The intracellular and extracellular compartments were neutralized and filled with either 150 mM KCl, NaCl or CsCl, and the extracellular compartment was filled with NaCl for all simulations. Hydrogen mass repartitioning (HMR) was applied using VMDs *hmr* plugin to enable a 4 fs time-step (**Fig. S1, Table S1**)^55^.

Simulations were conducted using GPU-accelerated NAMD 3.0^56^, CHARMM36m force-field was used for protein atoms, and CHARMM36 for lipids, ions and water^57^—TIP3P explicit water model. Initial energy minimization began with lipid melting simulations in the NVT ensemble, where non-lipid atoms where fixed and Langevin dynamics^58^ with a damping coefficient of 0.5 ps^–1^ applied for all atoms for 2 ns using a 1 fs timestep. This was followed by 2 ns of NPT equilibration with 1 kcal/mol restraints on protein atoms, using the Langevin Piston Nosé-Hoover^59^ method for pressure control at a 2 fs timestep. Restraints were then lifted from protein sidechains for and additional 2 ns with a 2 fs timestep. Each system was then subject to a 30 ns equilibration with all restraints removed, using a 4 fs timestep. Production simulations were run in replicate (n = 4) for 250 ns under constant volume (NVT) conditions with a 4 fs timestep, yielding a total of 1 µs of simulation time per condition. The final 200 ns of each short-timescale simulation were used for analysis.

### Constant electric-field simulations

Transjunctional voltage potentials (*V*_j_) were modeled using constant electric field (CEF) simulations, in which an external force was applied to all charged particles in the system^60,61^. The applied field strength was calculated according to:

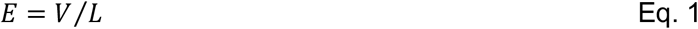

where *V* is the desired voltage and *L* is the height of the simulation-box (*i.e., c* or z-dimension). For a box height of 200 Å, field strengths corresponding to *V*_j_ values of: 0, 75, 100, or 200 mV were used. Systems with *V*_j_ values of ±150 mV were conducted under long-timescales (see below). In NAMD, voltages are defined in units of kcal/mol/Å/e, such that 0.0086, 0.012, and 0.023 correspond to 75, 100, and 200 mV, respectively. For the heterotypic Cx46/50 system, additional simulations were performed at -75 mV, -100 mV, and -200 mV. All CEF simulations reached steady state within approximately 30 ns, as assessed by monitoring the cation flux behavior.

### Long-timescale simulations

Long-timescale simulations of Cx46, Cx50, and heterotypic Cx46/50 channels were conducted on both Anton2^62^ and JURECA supercomputing platforms (see **Table S1**). Systems were prepared similarly to those described above, with simulations performed in orthorhombic boxes of dimensions 150×150×200 Å (Anton2) or 140×140×220 Å (JURECA) (**Fig. S1**). All systems were solvated with 150 mM KCl on the intracellular compartments and 150 mM NaCl extracellularly, and all atoms were described using the CHARMM36 force field, except for protein atoms, for which CHARMM36m was used.

For Anton2 simulations, systems were equilibrated locally using the protocol described for short-timescale simulations. The final frame of the 30 ns equilibration was used to initiate production runs. Simulations were performed in the NVT ensemble using Langevin dynamics with a 2.5 fs timestep. Hydrogen mass repartitioning was not applied. Each system was simulated in triplicate for ∼2 µs with an applied constant electric field of 0.0173 kcal/mol/Å/e, corresponding to a *V*_j_ of 150 mV (see Constant Electric-Field Simulations).

Simulations with JURECA were performed using Gromacs (version 2022)^63^, and the same forcefield (CHARMM36m for protein, and CHARMM36 for water, lipids and ions) was used for all simulations. V-rescale^64^ and C-rescale^65^ were used for temperature and pressure coupling, respectively. Equilibration protocols on JURECA followed the same scheme as those used for NAMD and Anton2. For long-timescale simulations, 250 ns blocks were used for certain analyses.

### Kinetics and thermodynamics of ion permeation

Average ionic currents (〈*I*〉) were calculated as the total number of permeation events (*N_p_*), over the total simulation time (*t*), multiplied by the ion valence (q):

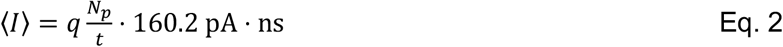

Currents were calculated as the slope from a linear fit to the cumulative ion counts over time^20,35^. A permeation event was defined as an ion sequentially moving from one bulk intracellular compartment (-90 < z < -60), through the pore region (|z| < 60), and into the opposite bulk compartment (60 < z < 90), or in reverse for a negative flux event. Junctional conductance was calculated from the slope of the current-voltage relationship defined by the (*I/V*) curve.

To analyze ion permeation energetics, one-dimensional Markov-state-models (MSMs) were constructed from the z-coordinates of permeating ions. The total z-length of the box was divided into 3 Å bins to define a discrete reaction coordinate. Bin-to-bin transitions were counted and row-normalized to generate a transition matrix (*T*), where each element (*T_i_*_,j_) describes the probability of transitioning from state *i* to state *j*, evaluated at a lag-time of 0.1 ns.

Under equilibrium conditions, the potential of mean force (PMF) was derived from the MSM transition matrix using the principle of detailed balance, as previously described^7,28^:

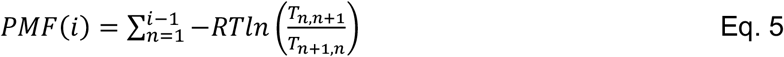

In non-equilibrium CEF simulations, the non-equilibrium driving potential (NEDP, *χ*) was calculated to describe the net force experienced by ions in steady-state flux conditions^66^:

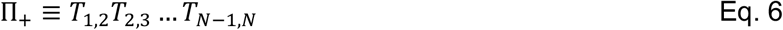

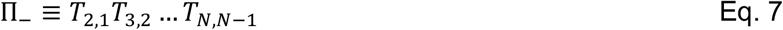

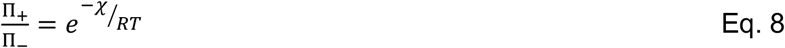

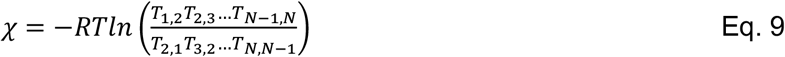

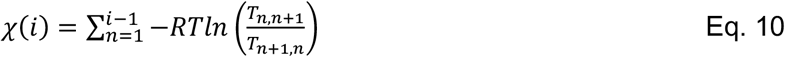

Here, Π_+_and Π_–_ are the cumulative product of the forward and reverse rate-constants within the MSM, respectively, and *RT* is the molar energy constant (kcal/mol). Rearrangement of Eq. 9 yields a function for *χ* at any state (i), which becomes mathematically equivalent to the PMF (Eq. 5) in the absence of external driving forces (where *χ* = 0).

### Thermodynamics and kinetics of K^+^ binding

Ion binding sites were identified by analyzing cation trajectories (K^+^, Cs^+^ or Na^+^) across short-timescale simulations. Ion-protein contact frequencies were determined by counting the number of ions within 3 Å of any protein atom in each trajectory frame (sampled every 100 ps). Additionally, time-averaged ion densities were calculated using the *volmap* plugin in VMD, based on the last 50 ns of each trajectory and averaged across four independent replicates. This approach revealed multiple discrete cation binding sites within the gap junction channels, designated as binding sites I–IV (see Results).

To quantify the thermodynamics of K^+^ binding at these sites, a two-state binding model was applied. Binding and unbinding events were tallied for each site in a protein-centric manner, disregarding the identity of individual ions and instead focusing on the occupancy state of the protein-defined site. Due to the six-fold rotational symmetry of the dodecameric gap junction channel, subunits were indexed such that chain i-1 is one subunit counterclockwise from chain i, and chain i+1 is one subunit clockwise relative to chain i. This indexing enabled the proper identification of interfacial binding sites formed by adjacent subunits.

The following selection strings were used to define binding sites I–IV (BS-I to BS-IV) within the connexin pore based on spatial proximity and side chain chemistry, and account for both intra- and inter-subunit contributions to ion coordination:

- **BS-I:** “protein and (resid 38 and name O and chain i)”
- **BS-II:** “protein and ((resid 49 50 52 and name O and chain i) or (resid 62 and name CD and chain i))”
- **BS-III:** “protein and ((((resid 54 56 and name O) or (resid 53 and name OG1) or (resid 62 and name OE1 OE2)) and chain i))”
- **BS-IV:** “protein and ((resid 48 and name OD1 OD2 and chain i) or (resid 47 51 and name OD1 OD2 and chain i+1))”

For each binding site, a minimum-distance trajectory was generated by tracking the nearest ion to each binding site at each time point of the trajectory. The resulting radial distance distributions (*i.e.,* minimum-distance) were used to define a Schmitt-Trigger filter, where the ‘lower-bound’ corresponds to the first minimum after the primary binding peak, and the ‘upper-bound’ was set at a maximum of 1 Å beyond the lower-bound. Binding-sites ‘II’ and ‘III’ had upper-bounds set 0.5 Å and 0.8 Å, respectively, beyond the lower-bound to ensure proximal binding-sites are not included in the ‘bound-state’ (see **Fig. S4**).

Minimum-distance trajectories were then converted into binary time series using the Schmitt-Trigger criteria, such that distances below the ‘lower-bound’ are considered bound (1), distances greater than the ‘upper-bound’ are considered unbound (0), and distances within the Schmitt-Trigger take the value of its most-recent state. This binary time series enabled identification of continuous bound and unbound intervals, from which dwell times were computed.

### Long-timescale dynamics and ICL-NT interactions

Computational single channel ion currents from long-timescale simulations were calculated using instantaneous current values (*I*(*t*)) defined as the number of permeations in a fixed time interval,

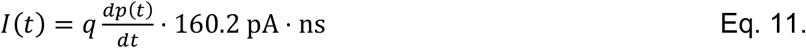

where *dp*(*t*) are the interpolated permeation events in a fixed time, *dt* = 0.1 ns, *q* is the ion valence, and *I*(*t*) is scaled by the conversion factor to yield pA. To reduce high-frequency noise and capture trends in dynamic current behavior, a uniform 100 ns running window average was applied across each interpolated current-trace. Minimum pore-radius (r_min_) values, calculated using the program HOLE2^67,68^, were defined as the minimum radius within each respective hemichannel (z = ±0–70 Å). These values were evaluated every 10 ns to monitor pore constriction events associated with conductance changes—similarly to reduce high-frequency noise, a uniform 50 ns running window average was performed.

To assess conductance states, ion current histograms were used to cluster trajectories into ‘low’, ‘mid’, and ‘high’ conductance regimes. Representative structures from each state were identified based on minimal root-mean-square-deviation (r.m.s.d.) to the average structure within the corresponding cluster. Interactions between the ICL and NT were quantified by generating pairwise distance matrices between the ICL region (residues 99–109), and NT domain (residues 2–15). A contact was defined as any residue pair between the two regions within 4 Å at a given frame. The frequency of contacts over time was used to assess the prevalence and dynamics of ICL- NT interactions across conductance states.

### Statistical analysis

All statistical comparisons are presented as the average plus or minus the standard error of the mean (s.e.m.) with n = 4 for short-timescale simulations and n = 3 for long-timescale simulations; p-values were evaluated using the non-parametric Mann-Whitney-U test for comparisons of distributions, the one-sided Student’s T-test was used to evaluate P_K+_/P_Cl–_ significance.

### Scripting and data visualization

All data analyses were performed in jupyter notebooks with Python3 the following relevant libraries: MDAnalysis (v2.5)^69,70^, NumPy (v1.26.4)^71^, Scikit-learn (v1.5.1)^72^, and SciPy (v1.13.1)^73^. Additional in-house scripts were developed for data preparation and analysis. All structural images and movies were rendered in ChimeraX (v1.16)^74^.

### Data and code availability

Initial structures and relevant codes used for thermodynamic calculations (*i.e.,* PMF, NEDP), ion tracking, and ICL-NT interactions are freely available at https://jugit.fz-juelich.de/computational-neurophysiology/connexin-46-50_conductance. Raw trajectory files will be available upon reasonable request to Bassam G. Haddad.

## ACKNOWLEDGMENTS

This work was supported by the National Institutes of Health (R35-GM124779 to S.L.R.) (R01-GM115805 to D.M.Z.) (F31-EY031580 to B.G.H.) and the National Science Foundation (MCB 1715823 to D.M.Z.). The authors gratefully acknowledge the computing time and staff support provided on Anton2 at PSC (MCB200098P to B.G.H.), Advanced Computing Center at Oregon Health & Science University (NIH S10OD034224). The authors gratefully acknowledge computing time on the supercomputer JURECA at Forschungszentrum Jülich (jara0177). Furthermore, the authors would like to acknowledge Prof. Dr. Jan-Philipp Machtens for his continued support.

## Author Contributions

B.G.H., D.M.Z., and S.L.R. designed, and planned the experiments. B.G.H. conducted and analyzed the data. B.G.H. and S.L.R. wrote the manuscript, and all authors contributed to final revisions. S.L.R. provided overall guidance to the design and execution of the work.

**Competing Interests:** The authors declare no competing interests.

## SUPPLEMENTAL MOVIE LEGENDS

`o` **Supplemental Movie S1. Representative K^+^ ion trajectory for Cx46.**
`o` **Supplemental Movie S2. Representative ‘fast’ K^+^ ion trajectory for Cx50.**
`o` **Supplemental Movie S3. Representative Cl^−^ ion trajectory for Cx50.**
`o` **Supplemental Movie S4. Representative current attenuation by ICL-NT interactions.**

## SUPPLEMENTAL FIGURES & LEGENDS

**Supplemental Figure S1.**
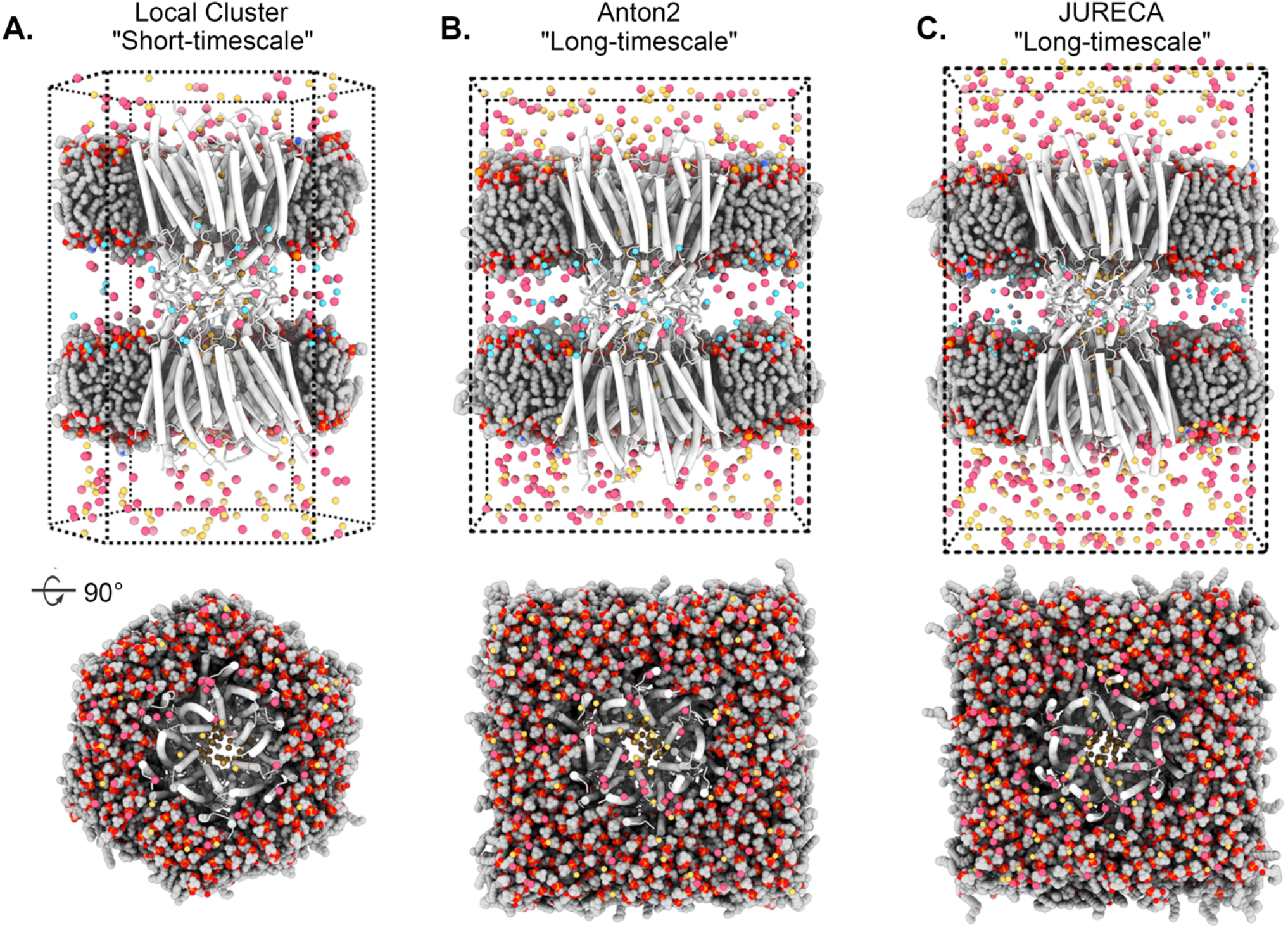
Depiction of MD simulation systems. **A**) System used for short-timescale simulations (4x 250 ns per condition) run on Exacloud (Oregon Health & Science University: Advanced Computing Center) using hexagonal periodic boundary conditions. **B**) System used for long-timescale simulations (3x ∼2 µs per condition), run on Anton2 (University of Pittsburgh: Pittsburgh Supercomputing Center). **C**) System for long-timescale simulations (5x ∼1–2 µs per condition), run on JURECA (Forschungszentrum Jülich: Jülich Supercomputing Center). All systems depict Cx46 (white) embedded in dual POPC bilayers (grey and red). Ions colored as K^+^ (yellow), Cl^−^ (red) and Na^+^ (blue). Explicit water molecules (TIP3) omitted for clarity.

**Supplemental Figure S2.**
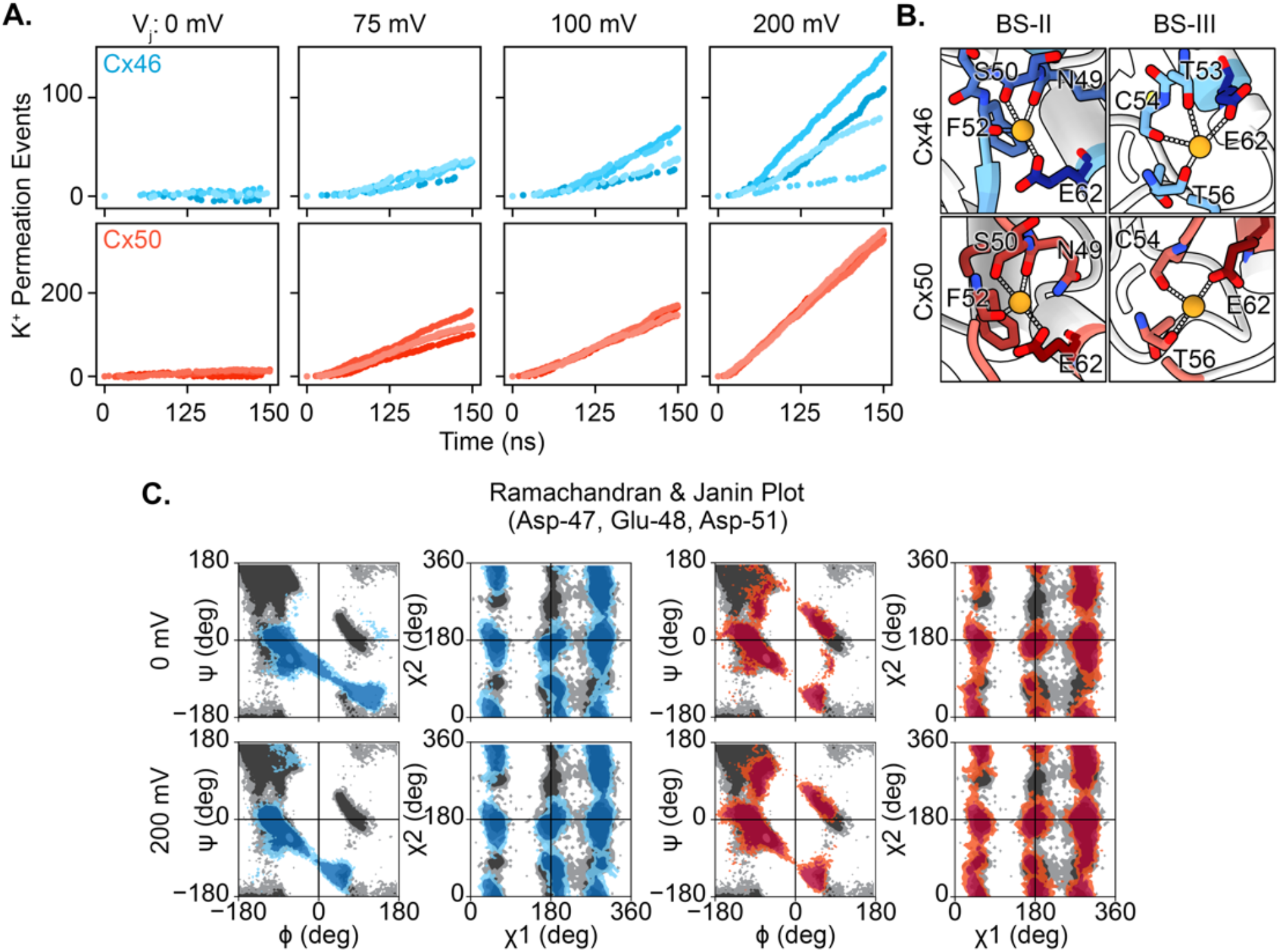
Supplemental characterization of K+ ion flux and binding sites for Cx46 and Cx50 simulations. **A**) Individual K^+^ flux traces for Cx46 (top) and Cx50 (bottom) at *V*_j_ = 0, 75, 100, and 200 mV. **B**) Representative visualizations of K^+^ binding sites II and III (BS-II and BS-III), shared between Cx46 (top) and Cx50 (bottom). In Cx46, T53 additionally contributes to K^+^ ion coordination for binding site III. **C**) Ramachandran and Janin plots for K^+^ binding site IV (BS-IV), formed by residues D47, E48, and D51, in Cx46 (left, blue) and Cx50 (right, red) at *V*_j_ = 0 mV (top) and 200 mV (bottom). Conformational differences at this site are expected to influence K^+^ binding interactions.

**Supplemental Figure S3.**
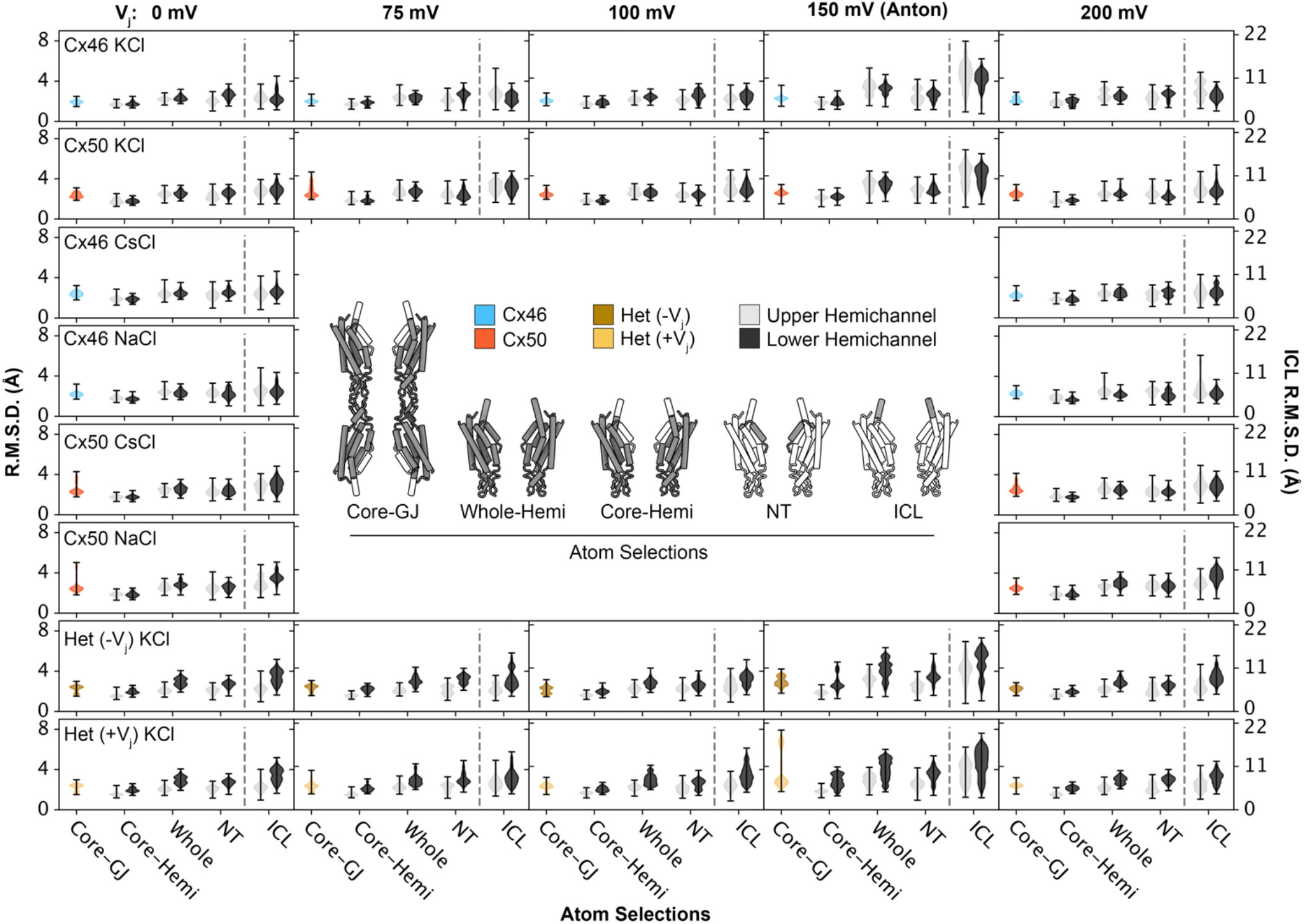
Root-mean-squared-deviation (RMSD) analysis for Cx46, Cx50 and heterotypic Cx46/50. Violin plots display RMSDs for Cx46, Cx50 and heterotypic Cx46/50 channels under varying internal ion conditions and *V*_j_ values (indicated). RMSD values are calculated for different structural regions (illustrated atom selections in dark gray) corresponding to: the core gap junction (core-GJ), which excludes the more dynamic NT and ICL regions; the whole hemichannel (Whole-Hemi); the core hemichannel (Core-Hemi); the N- terminal (NT) domain (residues 2-15); and the intracellular loop (ICL) region (residues 99-109). Values are shown for Cx50 (red), Cx46 (blue) and heterotypic Cx46/50 under -*V*_j_ (brown) and +*V*_j_ (yellow). RMSD values for Upper and Lower hemichannels are displayed in light gray and dark gray, respectively. Due to enhanced dynamics of the ICL region, its RMSD values are plotted on a separate right axis, while RMSD values for all other atom selections are shown on the left axis.

**Supplemental Figure S4.**
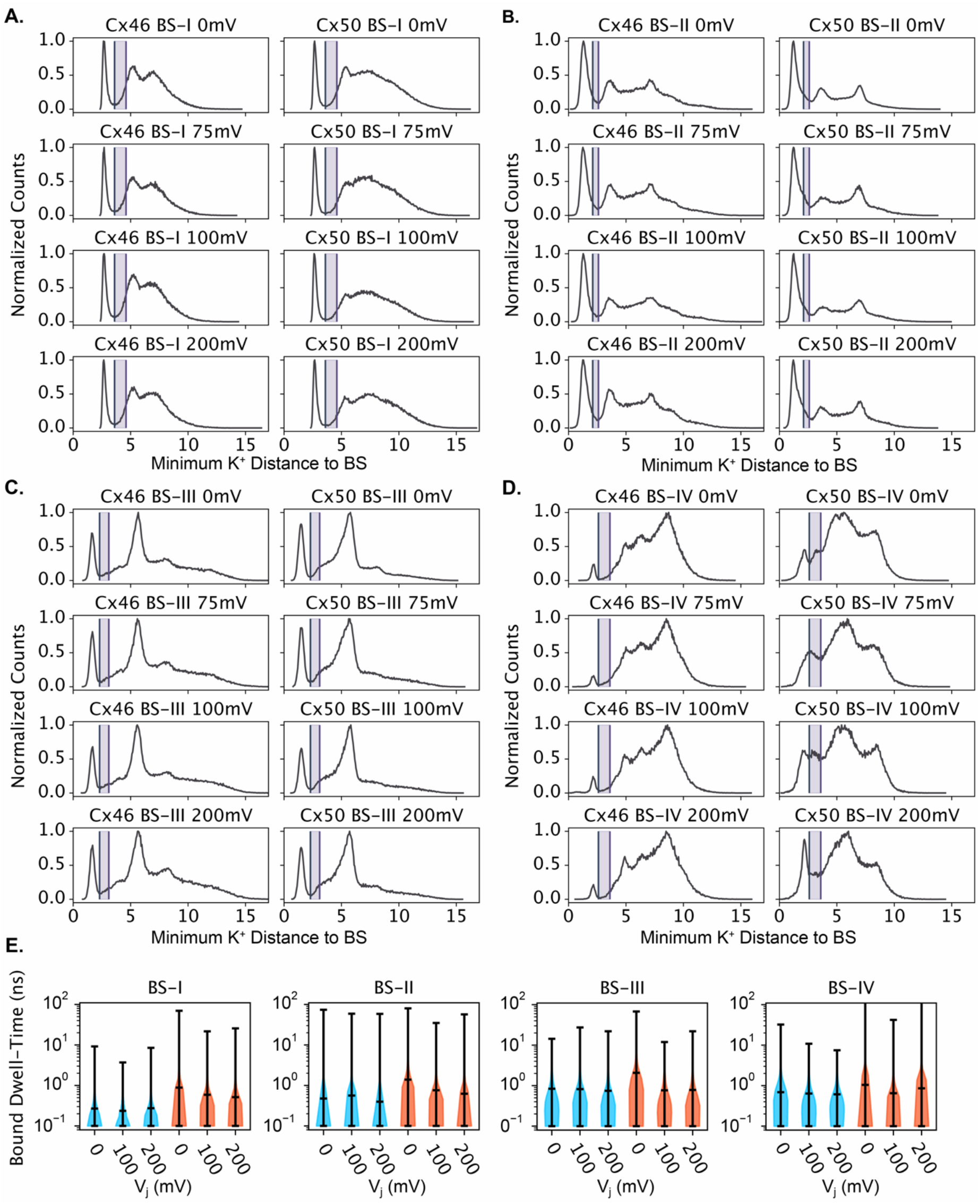
**Thermodynamic characterization of K+ binding sites. A–D**) Distance distributions of the nearest K^+^ ion to binding sites I-IV (BS-I to BS-IV) in Cx46 and Cx50 under *V*_j_ = 0, 75, 100 and 200 mV. Shaded regions denote the Schmitt-Trigger boundaries, which define bound and unbound states: distances left of the trigger are categorized as “Bound”, while those right of the trigger are “Unbound”. Ion positions within the trigger region are assigned based on their previous state (see Methods). **E**) Violin plots depicting bound dwell time (ns) for BS-I to BS-IV in Cx46 (blue) and Cx50 (red) at varying *V*_j_ potentials.

**Supplemental Figure S5.**
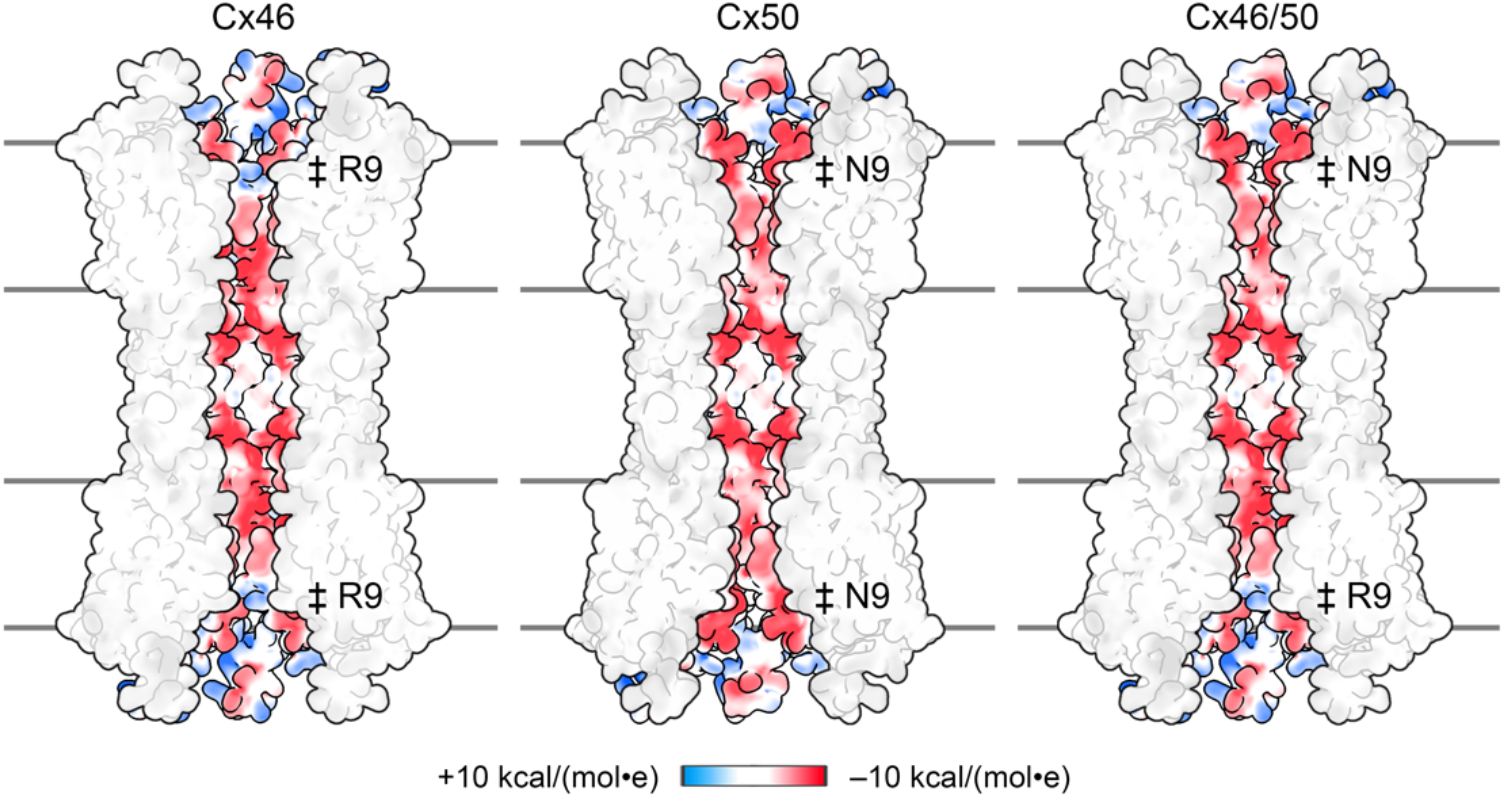
Pore electrostatic potential maps. Coulombic surface potential of Cx46 (left), Cx50 (middle) and heterotypic Cx46/50 (right) channels, shown in split-view to highlight the electrostatic properties of the permeation pathway. Coulombic surface-potential was calculated using ChimeraX’s couloumb-calculator, and displayed as positive (blue), neutral (white), and negative (red). Amino acid positions of R9 (Cx46) and N9 (Cx50) indicated (‡).

**Supplemental Figure S6.**
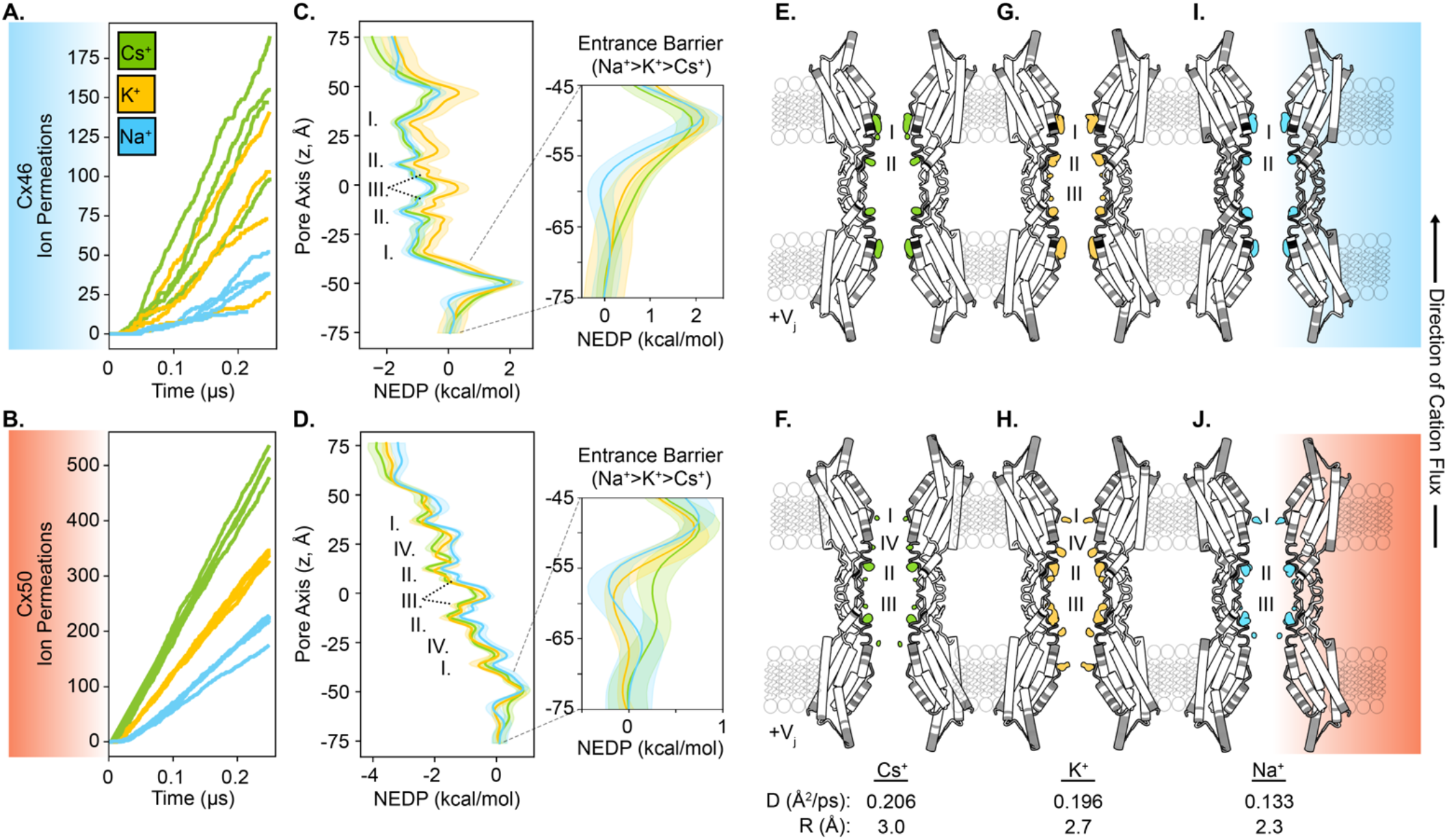
**Permeation of monovalent cations (Cs+, K+, and Na+) through homotypic Cx46 and Cx50 gap junctions. A,B**) Ion flux traces from simulations with intracellular solutions of Cs^+^ (green), K^+^ (yellow), Na^+^ (blue) for Cx46 (A) and Cx50 (B). **C,D**) Non-equilibrium driving potentials (NEDPs) for cation permeation at *V*_j_ = 200 mV for Cx46 (C) and Cx50 (D), displayed as mean ± s.e.m. calculated from multiple replicates (n=4). Interaction sites I–III (Cx46) and I–IV (Cx50) are annotated. Insets, show zoom-views of energetic entrance barriers (z = -75 to -45 Å). For Cx46, the Cs^+^ entrance barrier (1.85 +/- 0.23 kcal/mol) is approximately the same as that of K^+^ (2.07 +/- 0.36 kcal/mol; p = 0.67). Alternatively, in Cx50 the Cs^+^ entrance barrier (0.50 +/- 0.17 kcal/mol) is significantly lower than that of K^+^ (0.86 +/- 0.13 kcal/mol; p = 0.002). These differences suggest a rationale for the enhanced conductance of Cs^+^ over K^+^, which cannot be solely explained by differences in diffusion coefficients (D_Cs_/D_K_ = 1.05). **E-J**). Time-averaged cation density maps shown at equivalent iso-thresholds, colored by cation type, with binding sites labeled. Protein structures are shaded gray according to cation interaction frequency. Diffusion coefficients (D) and ionic radii (R) are indicated (bottom)^43^.

**Supplemental Figure 7.**
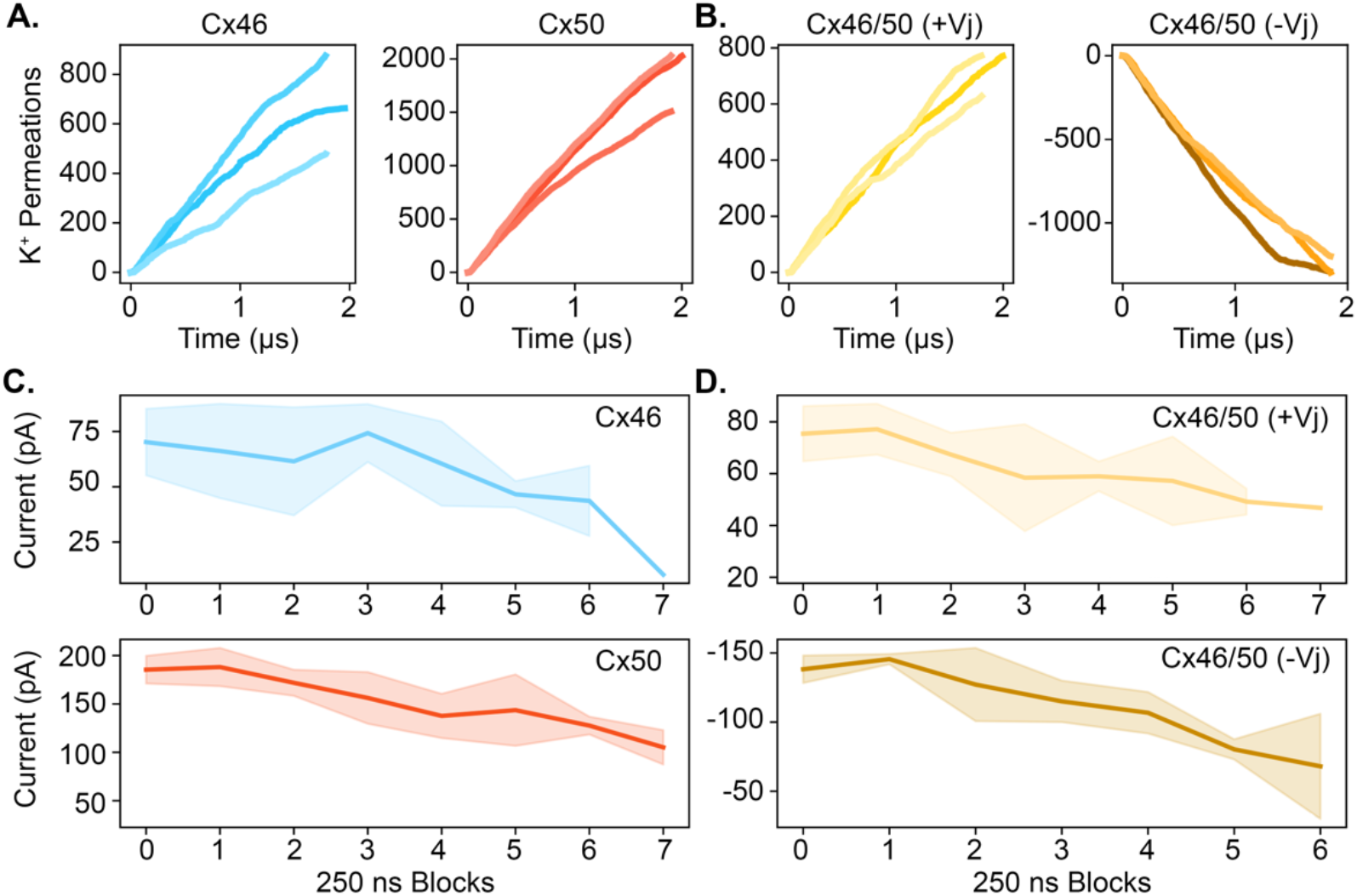
C**u**rrent **attenuation observed during long-timescale simulations of Cx46, Cx50, and Cx46-50 heterotypic gap junctions. A)** Cumulative K^+^ flux traces for Cx46 (blue) Cx50 (red) and **B**) heterotypic Cx46/50 at +*V*_j_ (yellow) and -*V*_j_ (brown). Data in A and B were obtained from long-timescale simulations at *V*_j_ = 150 mV. **C,D)** Block-averaged K^+^ current over time from long-timescale simulations, for Cx46 and Cx50 (C) and heterotypic Cx46/50 (D). Block sizes are 250 ns, and current values displayed as mean *±* s.d. (n=3), with shaded regions representing standard deviation. Regions without shading indicated time points where only one replicate contributed to data.

**Supplemental Figure S8.**
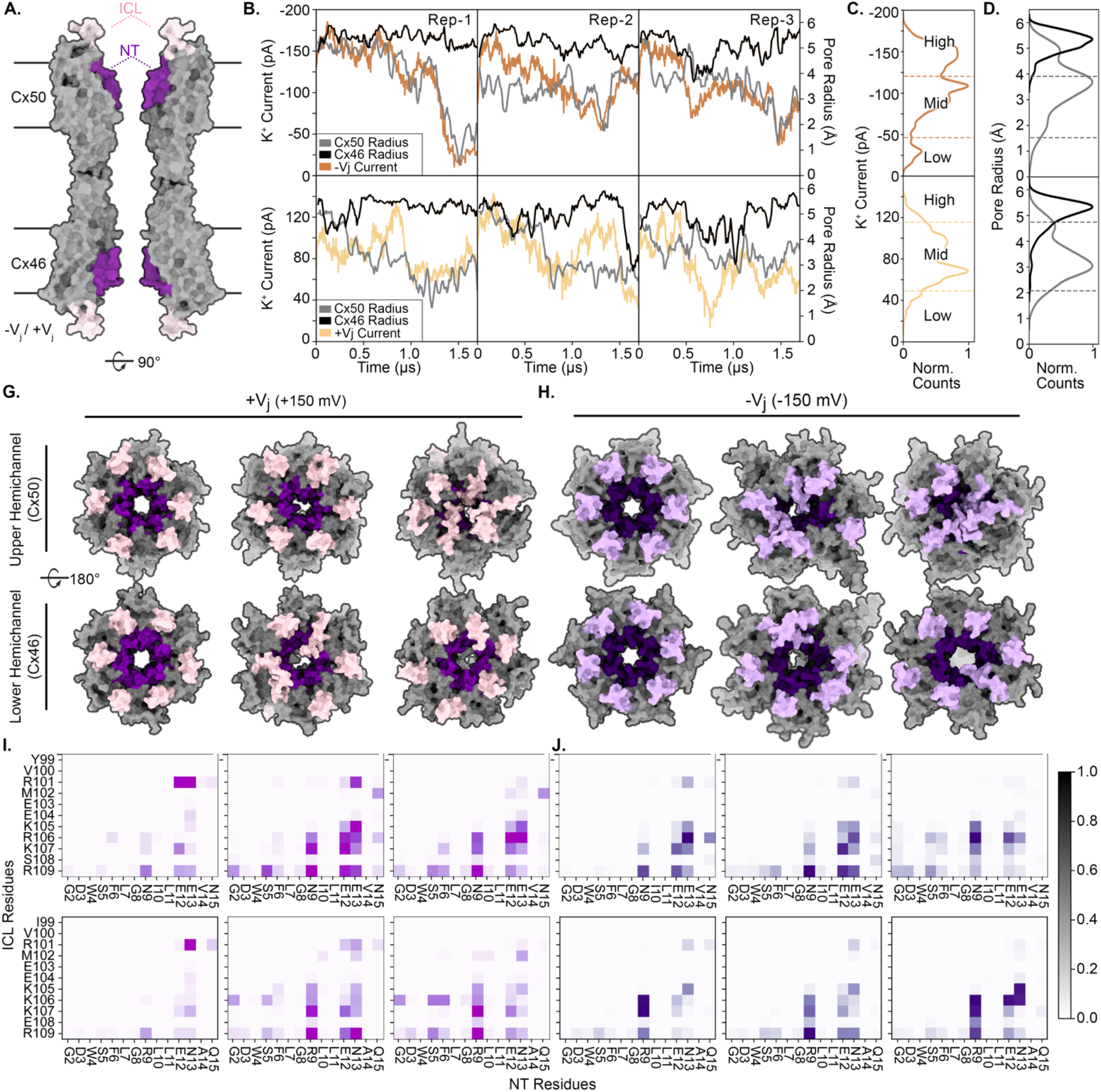
Attenuation of ion currents by dynamic ICL–NT interactions in heterotypic Cx46/50 gap junctions. **A**) Cross-sectional view of heterotypic Cx46/50 gap junction in surface representation, highlighting the NT (dark coloring) and ICL (light coloring). **B**) Traces of K^+^ current (left axis) and minimum pore radius (right axis) over time for Anton2 replicates (n=3; Rep-1 to Rep-3). Window-averaged K^+^ currents (100 ns windows) are shown in color, with -*V*_j_ (brown) and +*V*_j_ (yellow). Pore radius values for the Upper hemichannel (Upper; Cx50) and Lower hemichannel (Lower; Cx46) shown in gray and black, respectively. **C**) Distribution of K^+^ currents over time, clustered as ‘High, ‘Mid’, and ‘Low’ conductance states (dashed lines). **D**) Distributions of ‘Upper (Cx50)’ and ‘Lower (Cx46)’ pore radii, corresponding to the conductance-state clusters in Panel C (dashed lines). **E,F**) Views of the channel pore for mean structure for each conductance state in heterotypic Cx46/50 under +*V*_j_ (E) and -*V*_j_ (F). The Upper hemichannel (Cx50, top), and Lower hemichannel (Cx46, bottom) are shown separately. **G,H**) ICL–NT contact maps for the Upper (Cx50, top) and Lower (Cx46, bottom) hemichannels, corresponding to each conductance state under +*V*_j_ (G) and -*V*_j_ (H). Color intensity represents the normalized interaction frequency.

**Supplemental Figure S9.**
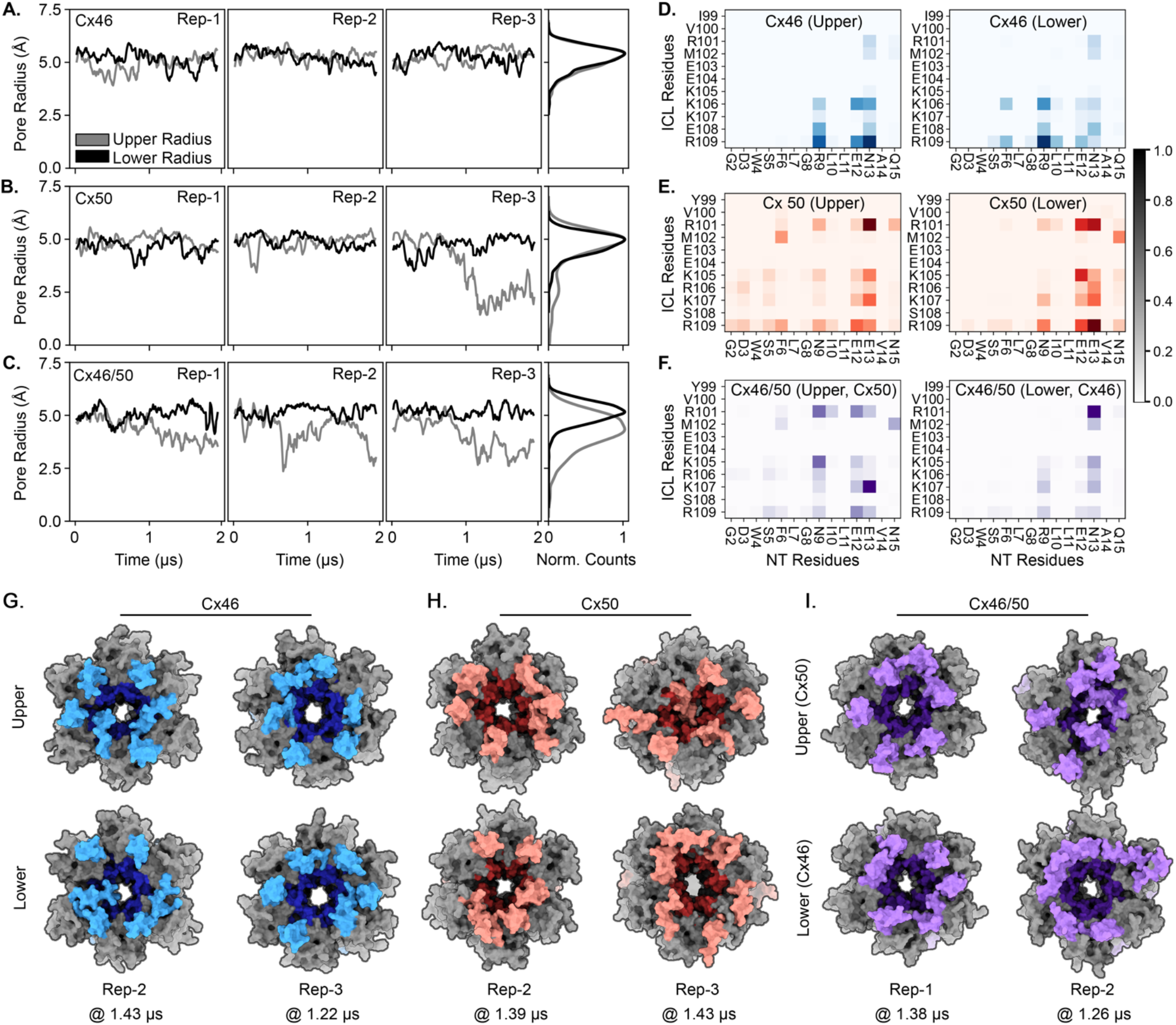
Pore profile analysis in the absence of Vj from long-timescale simulations. A–. **C**) Traces of minimum pore radius for Upper hemichannel (grey) and Lower hemichannel (black) over time, obtained for Cx46 (A), Cx50 (B), and the heterotypic Cx46/50 (C). Data are shown for three independent replicates (n=3; Rep-1 to Rep-3). Normalized pore radius distributions are displayed on the right. **D–F**) ICL–NT contact maps for the Upper (left) and Lower (right) hemichannels in Cx46 (D), Cx50 (E), and heterotypic Cx46/50 (F). Color intensity represents the normalized interaction frequency. Contact maps were generated from the last microsecond of each simulation, as minimal interactions were observed during the initial phase. **G–I**) Views of the channel pore depicting the mean structure for each system at different time points (indicated) representing open or partially occluded states, influenced by ICL (light shading) and NT (dark shading) interactions. Upper (top row) and Lower (bottom row) hemichannels are shown separately.

**Supplemental Figure S10.**
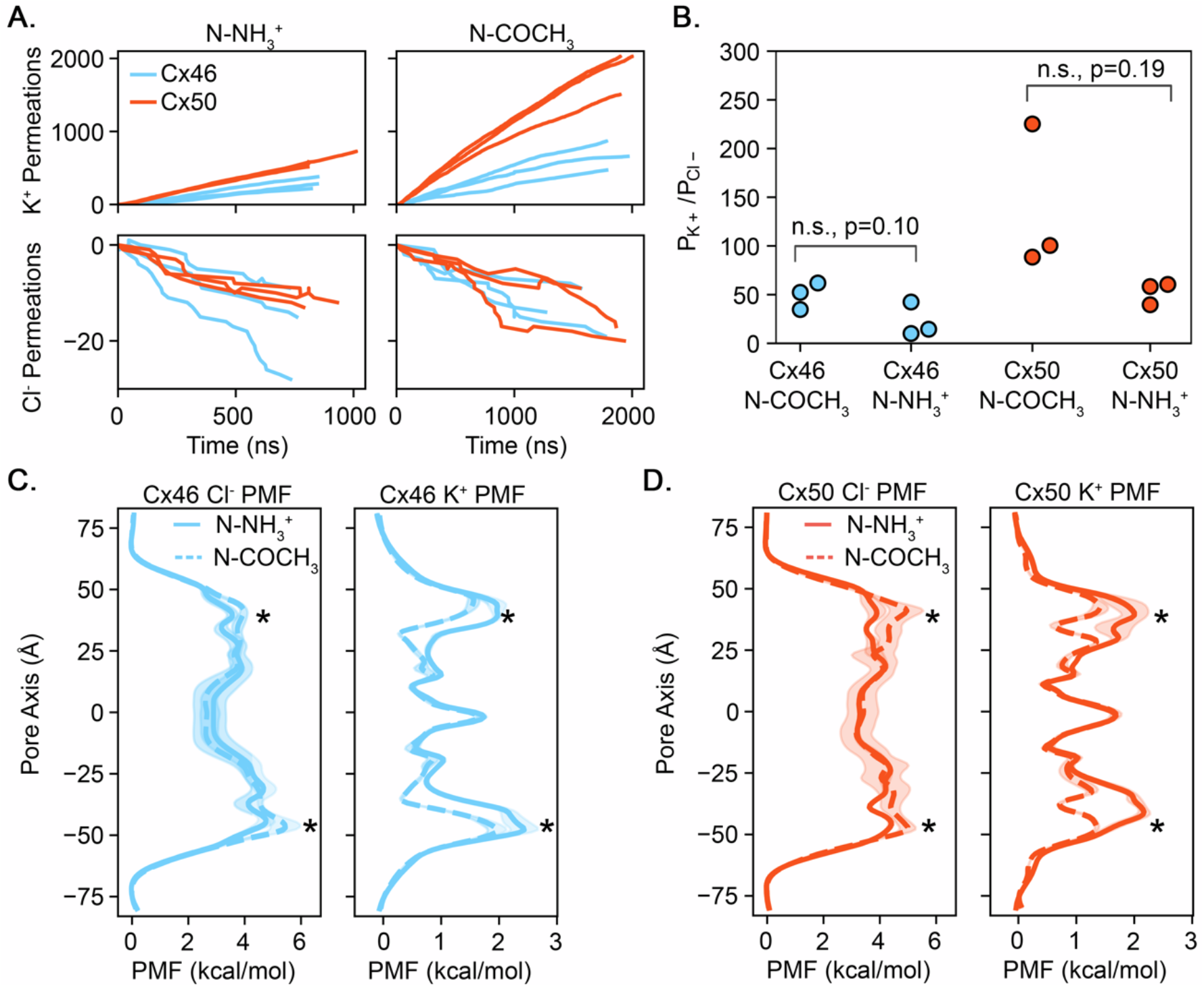
Effect of N-terminal acetylation on K+/Cl– permeation and selectivity. **A**) Cumulative K^+^ (top) and Cl^−^ (bottom) permeation events for Cx46 (blue) and Cx50 (red) simulated with an unacetylated N-terminus (N–NH_3_^+^; left) or acetylated N-terminus (N–COCH_3_; right). Each trace represents an independent simulation replicate. **B**) K⁺/Cl⁻ selectivity ratios (P_K⁺_/P_Cl⁻_) calculated from ion current ratios for acetylated and unacetylated channels. Differences did not reach statistical significance for either Cx46 or Cx50. **C–D**) Potential of mean force (PMF) profiles for Cl^−^ (left) and K^+^ (right) permeation through Cx46 (C) and Cx50 (D) with an unacetylated N-terminus (solid lines) or acetylated N-terminus (dashed lines). N-terminal acetylation produces modest changes in the entrance barriers encountered by K^+^ and Cl^−^ (asterisks), consistent with a modest effect on overall ion selectivity.

**Supplemental Table S1.**
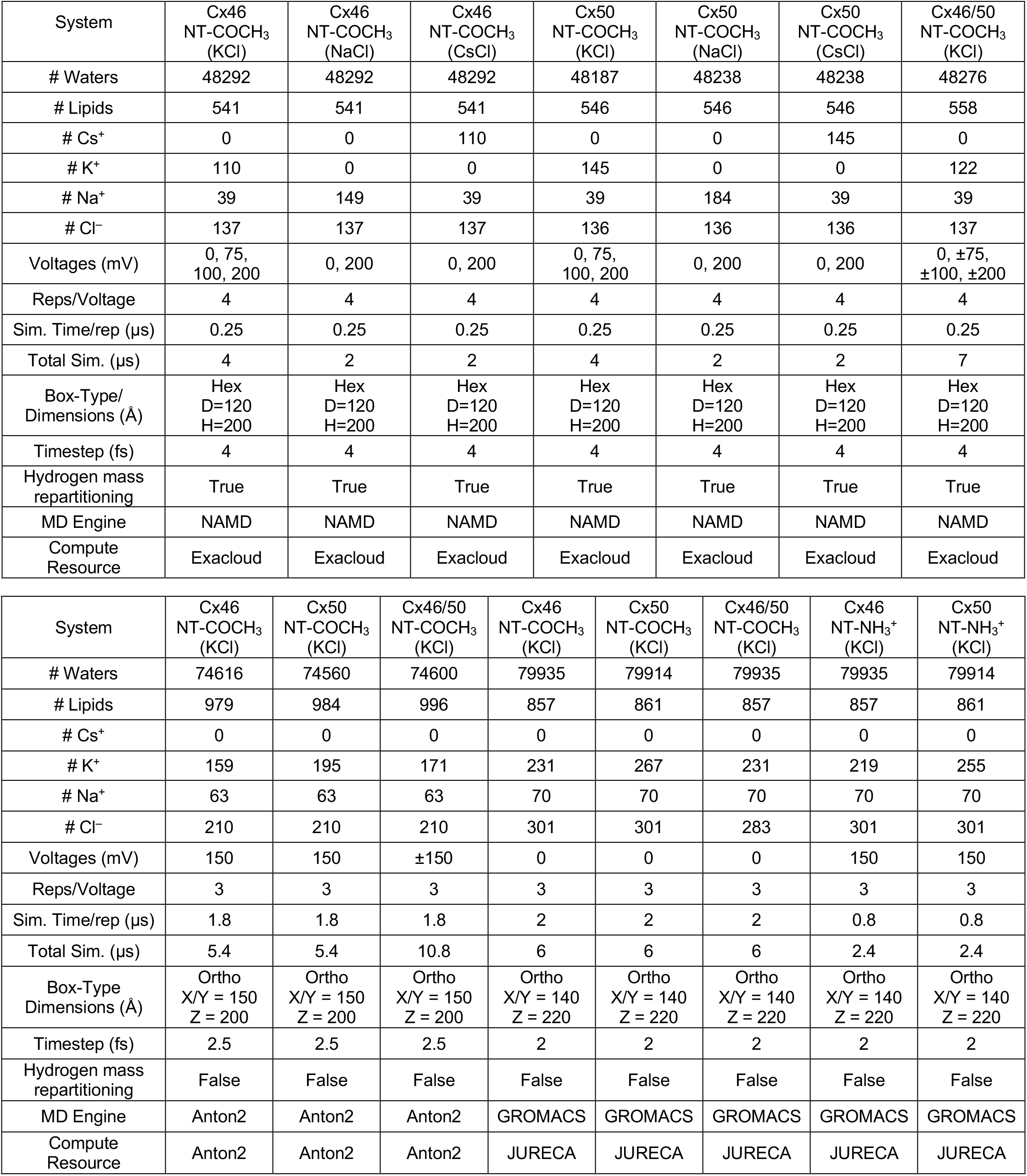
MD Simulation composition and set-up. Advanced compute-resources (Anton2, JURECA, and Exacloud) were used for production level simulations; equilibrations were performed on local workstations utilizing a range of consumer-grade NVIDIA GPUs.

| System | Cx46<br>NT-COCH<sub>3</sub><br>(KCl) | Cx46<br>NT-COCH<sub>3</sub><br>(NaCl) | Cx46<br>NT-COCH<sub>3</sub><br>(CsCl) | Cx50<br>NT-COCH<sub>3</sub><br>(KCl) | Cx50<br>NT-COCH<sub>3</sub><br>(NaCl) | Cx50<br>NT-COCH<sub>3</sub><br>(CsCl) | Cx46/50<br>NT-COCH<sub>3</sub><br>(KCl) |
| --- | --- | --- | --- | --- | --- | --- | --- |
| # Waters | 48292 | 48292 | 48292 | 48187 | 48238 | 48238 | 48276 |
| # Lipids | 541 | 541 | 541 | 546 | 546 | 546 | 558 |
| # Cs<sup>+</sup> | 0 | 0 | 110 | 0 | 0 | 145 | 0 |
| # K<sup>+</sup> | 110 | 0 | 0 | 145 | 0 | 0 | 122 |
| # Na<sup>+</sup> | 39 | 149 | 39 | 39 | 184 | 39 | 39 |
| # Cl<sup>-</sup> | 137 | 137 | 137 | 136 | 136 | 136 | 137 |
| Voltagess (mV) | 0, 75,<br>100, 200 | 0, 200 | 0, 200 | 0, 75,<br>100, 200 | 0, 200 | 0, 200 | 0, ±75,<br>±100, ±200 |
| Reps/Voltage | 4 | 4 | 4 | 4 | 4 | 4 | 4 |
| Sim. Time/rep (μs) | 0.25 | 0.25 | 0.25 | 0.25 | 0.25 | 0.25 | 0.25 |
| Total Sim. (μs) | 4 | 2 | 2 | 4 | 2 | 2 | 7 |
| Box-Type/<br>Dimensions (Å) | Hex<br>D=120<br>H=200 | Hex<br>D=120<br>H=200 | Hex<br>D=120<br>H=200 | Hex<br>D=120<br>H=200 | Hex<br>D=120<br>H=200 | Hex<br>D=120<br>H=200 | Hex<br>D=120<br>H=200 |
| Timestep (fs) | 4 | 4 | 4 | 4 | 4 | 4 | 4 |
| Hydrogen mass<br>repartitioning | True | True | True | True | True | True | True |
| MD Engine | NAMD | NAMD | NAMD | NAMD | NAMD | NAMD | NAMD |
| Compute<br>Resource | Exacloud | Exacloud | Exacloud | Exacloud | Exacloud | Exacloud | Exacloud |

| System | Cx46<br>NT-COCH<sub>3</sub><br>(KCl) | Cx50<br>NT-COCH<sub>3</sub><br>(KCl) | Cx46/50<br>NT-COCH<sub>3</sub><br>(KCl) | Cx46<br>NT-COCH<sub>3</sub><br>(KCl) | Cx50<br>NT-COCH<sub>3</sub><br>(KCl) | Cx46/50<br>NT-COCH<sub>3</sub><br>(KCl) | Cx46<br>NT-NH<sub>3</sub><sup>+</sup><br>(KCl) | Cx50<br>NT-NH<sub>3</sub><sup>+</sup><br>(KCl) |
| --- | --- | --- | --- | --- | --- | --- | --- | --- |
| # Waters | 74616 | 74560 | 74600 | 79935 | 79914 | 79935 | 79935 | 79914 |
| # Lipids | 979 | 984 | 996 | 857 | 861 | 857 | 857 | 861 |
| # Cs<sup>+</sup> | 0 | 0 | 0 | 0 | 0 | 0 | 0 | 0 |
| # K<sup>+</sup> | 159 | 195 | 171 | 231 | 267 | 231 | 219 | 255 |
| # Na<sup>+</sup> | 63 | 63 | 63 | 70 | 70 | 70 | 70 | 70 |
| # Cl<sup>-</sup> | 210 | 210 | 210 | 301 | 301 | 283 | 301 | 301 |
| Voltagess (mV) | 150 | 150 | ±150 | 0 | 0 | 0 | 150 | 150 |
| Reps/Voltage | 3 | 3 | 3 | 3 | 3 | 3 | 3 | 3 |
| Sim. Time/rep (μs) | 1.8 | 1.8 | 1.8 | 2 | 2 | 2 | 0.8 | 0.8 |
| Total Sim. (μs) | 5.4 | 5.4 | 10.8 | 6 | 6 | 6 | 2.4 | 2.4 |
| Box-Type<br>Dimensions (Å) | Ortho<br>X/Y = 150<br>Z = 200 | Ortho<br>X/Y = 150<br>Z = 200 | Ortho<br>X/Y = 150<br>Z = 200 | Ortho<br>X/Y = 140<br>Z = 220 | Ortho<br>X/Y = 140<br>Z = 220 | Ortho<br>X/Y = 140<br>Z = 220 | Ortho<br>X/Y = 140<br>Z = 220 | Ortho<br>X/Y = 140<br>Z = 220 |
| Timestep (fs) | 2.5 | 2.5 | 2.5 | 2 | 2 | 2 | 2 | 2 |
| Hydrogen mass<br>repartitioning | False | False | False | False | False | False | False | False |
| MD Engine | Anton2 | Anton2 | Anton2 | GROMACS | GROMACS | GROMACS | GROMACS | GROMACS |
| Compute<br>Resource | Anton2 | Anton2 | Anton2 | JURECA | JURECA | JURECA | JURECA | JURECA |

## REFERENCES

1 Bezanilla, F. Gating currents. J Gen Physiol 150, 911–932 (2018). 10.1085/jgp.201812090

2 González, C. et al. K(+) channels: function-structural overview. Compr Physiol 2, 2087–2149 (2012). 10.1002/cphy.c110047

3 Roux, B. Ion conduction and selectivity in K(+) channels. Annu Rev Biophys Biomol Struct 34, 153– 171 (2005). 10.1146/annurev.biophys.34.040204.144655

4 Bernèche, S. C Roux, B. Energetics of ion conduction through the K+ channel. Nature 414, 73–77 (2001). 10.1038/35102067

5 Zhang, K., Julius, D. & Cheng, Y. Structural snapshots of TRPV1 reveal mechanism of polymodal functionality. Cell 184, 5138–5150.e5112 (2021). 10.1016/j.cell.2021.08.012

6 Flores, J. A. et al. Connexin-46/50 in a dynamic lipid environment resolved by CryoEM at 1.9 A. Nat Commun 11, 4331 (2020). 10.1038/s41467-020-18120-5

7 Myers, J. B. et al. Structure of native lens connexin 46/50 intercellular channels by cryo-EM. Nature 564, 372–377 (2018). 10.1038/s41586-018-0786-7

8 Qi, C., et al. Structure of the connexin-43 gap junction channel in a putative closed state. Elife 12 (2023). 10.7554/eLife.87616

9 Lee, S. N. et al. Cryo-EM structures of human Cx36/GJD2 neuronal gap junction channel. Nat Commun 14, 1347 (2023). 10.1038/s41467-023-37040-8

10 Oshima, A. Structure of an innexin gap junction channel and cryo-EM sample preparation. Microscopy (Oxf) 66, 371–379 (2017). 10.1093/jmicro/dfx035

11 He, Z. et al. Structural and functional analysis of human pannexin 2 channel. Nat Commun 14, 1712 (2023). 10.1038/s41467-023-37413-z

12 Choi, W., Clemente, N., Sun, W., Du, J. & Lu, W. The structures and gating mechanism of human calcium homeostasis modulator 2. Nature 576, 163–167 (2019). 10.1038/s41586-019-1781-3

13 Rutz, S., Deneka, D., Dittmann, A., Sawicka, M. & Dutzler, R. Structure of a volume-regulated heteromeric LRRC8A/C channel. Nat Struct Mol Biol 30, 52–61 (2023). 10.1038/s41594-022-00899-0

14 Syrjanen, J., Michalski, K., Kawate, T. & Furukawa, H. On the molecular nature of large-pore channels. Journal of Molecular Biology 433, 166994 (2021). 10.1016/j.jmb.2021.166994

15 Goodenough, D. A. & Paul, D. L. Gap junctions. Cold Spring Harb Perspect Biol 1, a002576 (2009). 10.1101/cshperspect.a002576

16 Lucaciu, S. A., Leighton, S. E., Hauser, A., Yee, R. & Laird, D. W. Diversity in connexin biology. J Biol Chem 29, 105263 (2023). 10.1016/j.jbc.2023.105263

17. Abrams, C. K., et al. Properties of human connexin 31, which is implicated in hereditary dermatological disease and deafness. PNAS (2006).

18 Oh, S. et al. Changes in Permeability Caused by Connexin 32 Mutations Underlie X-Linked Charcot-Marie-Tooth Disease. Neuron (1997). 10.1016/s0896-6273(00)80973-3

19 Trexler, E. B., Bennett, M. V., Bargiello, T. A. & Verselis, V. K. Voltage gating and permeation in a gap junction hemichannel. Proceedings of the National Academy of Sciences 93, 5836–5841 (1996). doi:10.1073/pnas.93.12.5836

20 Harris, A. L. Connexin channel permeability to cytoplasmic molecules. Prog Biophys Mol Biol 94, 120–143 (2007). 10.1016/j.pbiomolbio.2007.03.011

21 Trexler, E. B., Bukauskas, F. F., Kronengold, J., Bargiello, T. A. & Verselis, V. K. The First Extracellular Loop Domain Is a Major Determinant of Charge Selectivity in Connexin46 Channels. Biophys J (2000).

22 Bukauskas, F. F. & Verselis, V. K. Gap junction channel gating. Biochimica et Biophysica Acta (BBA) - Biomembranes 1662, 42–60 (2004). 10.1016/j.bbamem.2004.01.008

23 Xin, L., Gong, X.-Q. & Bai, D. The Role of Amino Terminus of Mouse Cx50 in Determining Transjunctional Voltage-Dependent Gating and Unitary Conductance. Biophysical Journal 99, 2077–2086 (2010). 10.1016/j.bpj.2010.07.032

24 Tong, J.-J. et al. Molecular mechanisms underlying enhanced hemichannel function of a cataract- associated Cx50 mutant. Biophysical Journal 120, 5644–5656 (2021). 10.1016/j.bpj.2021.11.004

25 Verselis, V. K., Ginter, C. S. & Bargiello, T. A. Opposite voltage gating polarities of two closely related onnexins. Nature 368, 348–351 (1994). 10.1038/368348a0

26 Kronengold, J., Srinivas, M. & Verselis, V. K. The N-Terminal Half of the Connexin Protein Contains the Core Elements of the Pore and Voltage Gates. The Journal of Membrane Biology 245, 453–463 (2012). 10.1007/s00232-012-9457-z

27 Ek Vitorín, José F., Pontifex, Tasha K. C Burt, Janis M. Determinants of Cx43 Channel Gating and Permeation: The Amino Terminus. Biophysical Journal 110, 127–140 (2016). 10.1016/j.bpj.2015.10.054

28 Yue, B. et al. Connexin 46 and connexin 50 gap junction channel properties are shaped by structural and dynamic features of their N-terminal domains. The Journal of Physiology 599, 3313–3335 (2021). 10.1113/JP281339

29 Jarodsky, J. M., Myers, J. B. & Reichow, S. L. Reversible lipid-mediated pH-gating of connexin- 46/50 by cryo-EM. Nature Communications 17, 1606 (2026). 10.1038/s41467-026-68311-9

30 Flores, J. A., O’Neill, S. E., Jarodsky, J. M. & Reichow, S. L. Calcium induced N-terminal gating and pore collapse in connexin-46/50 gap junctions. bioRxiv, 2025.2002.2012.637955 (2025). 10.1101/2025.02.12.637955

31 Peracchia, C. Chemical gating of gap junction channels: Roles of calcium, pH and calmodulin. Biochimica et Biophysica Acta (BBA) - Biomembranes 1662, 61–80 (2004). 10.1016/j.bbamem.2003.10.020

32 Maeda, S. et al. Structure of the connexin 26 gap junction channel at 3.5 A resolution. Nature 458, 597–602 (2009). 10.1038/nature07869

33 Wang, X., Li, L., Peracchia, L. L. & Peracchia, C. Chimeric evidence for a role of the connexin cytoplasmic loop in gap junction channel gating. Pflugers Arch 431, 844–852 (1996). 10.1007/s004240050076

34 Xu, Q., et al. Gating of connexin 43 gap junctions by a cytoplasmic loop calmodulin binding domain. Am J Physiol Cell Physiol 302, C1548–1556 (2012). 10.1152/ajpcell.00319.2011

35 Oh, S., Rubin, J. B., Bennett, M. V. L., Verselis, V. K. & Bargiello, T. A. Molecular Determinants of Electrical Rectification of Single Channel Conductance in Gap Junctions Formed by Connexins 26 and 32. J Gen Physiol (1999).

36 Rash, J. E. et al. Heterotypic gap junctions at glutamatergic mixed synapses are abundant in goldfish brain. Neuroscience 285, 166–193 (2015). 10.1016/j.neuroscience.2014.10.057

37 Hopperstad, M. G., Srinivas, M. & Spray, D. C. Properties of Gap Junction Channels Formed by Cx46 Alone and in Combination with Cx50. Biophysical Journal 79, 1954–1966 (2000). 10.1016/S0006-3495(00)76444-7

38 Zonta, F., Polles, G., Sanasi, M. F., Bortolozzi, M. & Mammano, F. The 3.5 ångström X−ray structure of the human connexin26 gap junction channel is unlikely that of a fully open channel. BMC Cell Biol (2013).

39 Srinivas, M., Kronengold, J., Bukauskas, F. F., Bargiello, T. A. & Verselis, V. K. Correlative studies of gating in Cx46 and Cx50 hemichannels and gap junction channels. Biophys J 88, 1725–1739 (2005). 10.1529/biophysj.104.054023

40 Bennett, B. C. et al. An electrostatic mechanism for Ca(2+)-mediated regulation of gap junction channels. Nat Commun 7, 8770 (2016). 10.1038/ncomms9770

41 Tong, X. et al. The First Extracellular Domain Plays an Important Role in Unitary Channel Conductance of Cx50 Gap Junction Channels. PLoS One 10, e0143876 (2015). 10.1371/journal.pone.0143876

42 Srinivas, M. et al. Voltage dependence of macroscopic and unitary currents of gap junction channels formed by mouse connexin50 expressed in rat neuroblastoma cells. The Journal of Physiology 517 (1999).

43 Duan, P. C., Le, X., Ashutosh, T., Gerhard, M. & Bob, E. Selectivity and Permeation in Calcium Release Channel of Cardiac Muscle: Alkali Metal Ions. Biophysical Journal 76, 1346–1366 (1999). 10.1016/S0006-3495(99)77297-8

44 Spray, D. C., Duffy, H. S. & Scemes, E.

45 Rubin, J. B., Verselis, V. K., Bennett, M. V. & Bargiello, T. A. Molecular analysis of voltage dependence of heterotypic gap junctions formed by connexins 26 and 32. Biophys J 62, 183–193; discussion 193–185 (1992). 10.1016/S0006-3495(92)81804-0

46 A.G. Volkov, S. P., D.W. Deamer,. Two mechanisms of permeation of small neutral molecules and hydrated ions across phospholipid bilayers. Bioelectrochemistry and Bioenergetics 42, 153–160 (1997). 10.1016/S0302-4598(96)05097-0.

47 Kwon, T., Harris, A. L., Rossi, A. & Bargiello, T. A. Molecular dynamics simulations of the Cx26 hemichannel: evaluation of structural models with Brownian dynamics. J Gen Physiol 138, 475– 493 (2011). 10.1085/jgp.201110679

48 Chen, H. & Bai, D. The rectification of heterotypic Cx46/Cx50 gap junction channels depends on intracellular magnesium. Biophysics Reports 10, 336–348 (2024). 10.52601/bpr.2024.240015

49 Valiunas, V., Brink, P. R. & White, T. W. Lens Connexin Channels Have Differential Permeability to the Second Messenger cAMP. Invest Ophthalmol Vis Sci 60, 3821–3829 (2019). 10.1167/iovs.19-27302

50 Jiang, W. et al. Free energy and kinetics of cAMP permeation through connexin26 via applied voltage and milestoning. Biophys J 120, 2969–2983 (2021). 10.1016/j.bpj.2021.06.024

51 Weber, P. A., Chang, H. C., Spaeth, K. E., Nitsche, J. M. C Nicholson, B. J. The permeability of gap junction channels to probes of different size is dependent on connexin composition and permeant-pore affinities. Biophys J 87, 958–973 (2004). 10.1529/biophysj.103.036350

52 Brink, P. R., Valiunas, V. & White, T. W. Lens Connexin Channels Show Differential Permeability to Signaling Molecules. International Journal of Molecular Sciences 21 (2020).

53 William, H., Andrew, D. & Klaus, S. VMD -- V isual M olecular D ynamics. Journal of Molecular Graphics 14, 33–38 (1996).

54 Jo, S., Kim, T., Iyer, V. G. & Im, W. CHARMM-GUI: A web-based graphical user interface for CHARMM. Journal of Computational Chemistry **2G**, 1859–1865 (2008). 10.1002/jcc.20945

55 Balusek, C. et al. Accelerating Membrane Simulations with Hydrogen Mass Repartitioning. Journal of Chemical Theory and Computation 15, 4673–4686 (2019). 10.1021/acs.jctc.9b00160

56 Phillips, J. C. et al. Scalable molecular dynamics on CPU and GPU architectures with NAMD. The Journal of Chemical Physics 153 (2020). 10.1063/5.0014475

57 Huang, J. & MacKerell, A. D., Jr. CHARMM36 all-atom additive protein force field: validation based on comparison to NMR data. J Comput Chem 34, 2135–2145 (2013). 10.1002/jcc.23354

58 Davidchack, R. L., Handel, R. & Tretyakov, M. V. Langevin thermostat for rigid body dynamics. J Chem Phys 130, 234101 (2009). 10.1063/1.3149788

59 Evans, D. J. & Holian, B. L. The Nose–Hoover thermostat. The Journal of Chemical Physics 83, 4069–4074 (1985). 10.1063/1.449071

60 Gumbart, J., Khalili-Araghi, F., Sotomayor, M. & Roux, B. Constant electric field simulations of the membrane potential illustrated with simple systems. Biochimica et Biophysica Acta (BBA) - Biomembranes 1818, 294–302 (2012). 10.1016/j.bbamem.2011.09.030

61 Roux, B. The Membrane Potential and its Representation by a Constant Electric Field in Computer Simulations. Biophysical Journal 95, 4205–4216 (2008). 10.1529/biophysj.108.136499

62 Shaw, D. E. et al. 41–53 (2014).

63 Mark James Abraham, T. M., Roland Schulz, Szilárd Páll, Jeremy C. Smith, Berk Hess, Erik Lindahl. GROMACS: High performance molecular simulations through multi-level parallelism from laptops to supercomputers. SoftwareX 1-2, 19–25 (2015). 10.1016/j.softx.2015.06.001

64 Bussi, G., Donadio, D. & Parrinello, M. Canonical sampling through velocity rescaling. J Chem Phys 126, 014101 (2007). 10.1063/1.2408420

65 Bernetti, M. C Bussi, G. Pressure control using stochastic cell rescaling. J Chem Phys 153, 114107 (2020). 10.1063/5.0020514

66 Hill, T. L. in Free Energy Transduction in Biology (ed Terrell L. Hill) 33–56 (Academic Press, 1977).

67 Smart, O. S., Goodfellow, J. M. & Wallace, B. A. The pore dimensions of gramicidin A. Biophys J 65, 2455–2460 (1993). 10.1016/s0006-3495(93)81293-1

68 Smart, O. S., Neduvelil, J. G., Wang, X., Wallace, B. A. & Sansom, M. S. P. HOLE: A program for the analysis of the pore dimensions of ion channel structural models. Journal of Molecular Graphics 14, 354–360 (1996). 10.1016/S0263-7855(97)00009-X

69 Michaud-Agrawal, N., Denning, E. J., Woolf, T. B. & Beckstein, O. MDAnalysis: A toolkit for the analysis of molecular dynamics simulations. Journal of Computational Chemistry 32, 2319–2327 (2011). 10.1002/jcc.21787

70 Gowers, R. J. et al.in *SciPy*.

71 Harris, C. R. et al. Array programming with NumPy. Nature 585, 357–362 (2020). 10.1038/s41586-020-2649-2

72 Pedregosa, F. et al. Scikit-learn: Machine Learning in Python. J. Mach. Learn. Res. 12, 2825–2830 (2011).

73 Virtanen, P. et al. SciPy 1.0: fundamental algorithms for scientific computing in Python. Nature Methods 17, 261–272 (2020). 10.1038/s41592-019-0686-2

74 Pettersen, E. F. et al. UCSF ChimeraX: Structure visualization for researchers, educators, and developers. Protein Science 30, 70–82 (2021). 10.1002/pro.3943

